# Integrative multi-omics increase resolution of the sea urchin posterior gut gene regulatory network at single cell level

**DOI:** 10.1101/2023.05.12.540495

**Authors:** Danila Voronov, Periklis Paganos, Marta S. Magri, Claudia Cuomo, Ignacio Maeso, Jose Luis Gómez-Skarmeta, Maria Ina Arnone

**Affiliations:** Stazione Zoologica Anton Dohrn, Naples, Italy; Centro Andaluz de Biología del Desarrollo, CSIC/Universidad Pablo de Olavide, Sevilla, Spain; University of Barcelona, Barcelona, Spain

**Keywords:** gene regulatory networks, sea urchin, embryo, gut, multi-omics, *Pdx1*

## Abstract

Drafting gene regulatory networks (GRNs) requires embryological knowledge pertaining to the cell type families, information on the regulatory genes, causal data from gene knockdown experiments and validations of the identified interactions by cis-regulatory analysis. We use multi-omics involving next-generation sequencing (-seq) to obtain the necessary information drafting the sea urchin posterior gut GRN. Here we present an update to the GRN using i) a single cell RNA-seq derived cell atlas highlighting the 2 day post fertilization (dpf) sea urchin gastrula cell type families, as well as the genes expressed at single cell level, ii) a set of putative cis-regulatory modules and transcription factor (TF) binding sites obtained from chromatin accessibility ATAC-seq data, and iii) interactions directionality obtained from differential bulk RNA-seq following knockdown of the TF Sp-Pdx1, a key regulator of gut patterning in sea urchins. Combining these datasets, we draft the GRN for the hindgut *Sp-Pdx1* positive cells in the 2 dpf gastrula embryo. Overall, our data resolves the complex connectivity of the posterior gut GRN and increases the resolution of gene regulatory cascades operating within it.

## Introduction

Gene regulatory networks (GRNs) are used to describe molecular underpinnings of developmental processes that lead to establishment of various cell and tissue types in a developing embryo. GRNs consist of two main components: genes, which are the nodes of the network, and their interactions, which are the edges of the network. Such networks allow visualization of inputs and outputs of transcription factors, signaling molecules and terminal differentiation genes in time and location specific manner.

Echinoderms, and in particular the purple sea urchin *Strongylocentrotus purpuratus,* have long been excellent experimental systems for evolutionary developmental studies (Arnone et al., 2015; McClay, 2011). Together with hemichordates, echinoderms form the ambulacraria group, that is a phylogenetic sister group to chordates (Röttinger and Lowe, 2012). GRNs were used to study and describe the molecular underpinnings of their embryonic development. The sea urchin endomesoderm GRN, in particular, is one of the best studied (Cary et al., 2020; Ettensohn, 2020; Massri et al., 2023; Peter and Davidson, 2010). In addition to that, the sea urchin embryonic gut GRN, which operates downstream of the endomesoderm one, from mid gastrula up until 3 days post fertilization (dpf) pluteus larva, has also been a subject of multiple studies, reviewed in Annunziata et al. (2019), showing the crucial role of two Parahox genes, *Sp-Pdx1* and *Sp-Cdx,* in the development of the sea urchin hindgut and pyloric sphincter.

The traditional protocol for drafting GRNs requires embryological knowledge pertaining to the cell and tissue types present in the embryo, knowledge of the regulatory genes describing the regulatory state of the tissue or cell type, causal information from perturbation experiments and validations of the GRNs through the cis-regulatory analysis, identifying the cis-regulatory modules (CRMs) of the regulatory genes (Materna and Oliveri, 2008). The process of identification of the cell types and expressed genes usually involves gene expression visualization techniques such as *in situ* hybridization, while identification of CRMs is usually achieved via sequence alignment with evolutionary closely related species (Lee et al., 2007; Livi and Davidson, 2007) to identify conserved regions that could play a role as CRMs. Such methods allow identification of only a subset of the expressed genes, usually highly expressed, for which *in situ* probes were made, and a limited number of CRMs around them. However, each cell expresses multiple genes that are controlled by various CRMs (Cui et al., 2017; de-Leon and Davidson, 2010; Paganos et al., 2021), thus high throughput approaches are essential to build complete GRNs.

With the advent of novel technologies involving next generation sequencing, such as Assay for Transposase-Accessible Chromatin with next-generation sequencing (ATAC-seq) (Buenrostro et al., 2015), bulk and single cell RNA sequencing (RNA-seq and scRNA-seq, respectively) high throughput identification of CRMs and transcription factors (TFs) bound to them (2018), gene expression profiles (Davidson et al., 2022; Massri et al., 2021; Paganos et al., 2021; Paganos et al., 2022c) and causal dynamics (Lowe et al., 2016; Rafiq et al., 2014) became possible. For example, Shashikant et al. (2018) used ATAC-seq to assess chromatin accessibility in sea urchin skeletogenic cells and were able to confirm previously known enhancers and identify more CRMs binding Sp-Alx1 and Sp-Ets1 TFs. Paganos et al. (2021) have used scRNA-seq data to identify the cell type families present in the 3 dpf sea urchin pluteus larva and to get the expression profiles of genes within every identified cell population at unprecedented resolution.

Here we present a cell type family atlas of the 2 dpf *S. purpuratus* gastrula obtained using single cell RNA-seq data and the set of putative gut CRMs at 2 dpf obtained with embryonic gut and whole embryo ATAC-seq, which we combine along with differential expression analyses to increase resolution of the hindgut GRN around *Sp-Pdx1* gene in *Sp-Pdx1* positive cells. We use ATAC-seq data for predicting CRMs and performing TF footprinting along with scRNA-seq data to narrow genome wide predictions to specific cell type families. Furthermore, we use the available bulk differential RNA-seq data to give causal information to the interactions within a GRN of the cells of the hindgut expressing the key gut patterning ParaHox gene *Sp-Pdx1* (Annunziata and Arnone, 2014; Cole et al., 2009). Thus, we use multi-omics datasets to obtain information necessary for GRN drafting as per established logic for drafting GRNs (Materna and Oliveri, 2008), and provide insight to whether the GRN interactions are predicted to be direct or not. Finally, using a combination of reporter gene mediated cis-regulatory analysis and TF knock-down in trans, we demonstrate *in vivo* validation of a predicted by our approach interaction between the Hox gene *Sp*-*Hox11/13b* and the ParaHox gene *Sp*-*Pdx1*.

## Results

### Chromatin accessibility predicts active CRMs in sea urchin gastrula nuclei

In order to identify the locations of putative CRMs (pCRMs) in the sea urchin gastrula and predict which TFs are bound to them, we have generated ATAC-seq libraries for the sea urchin gastrula whole embryo and the corresponding isolated gut tissue in two replicates each. The consensus set of peaks was obtained by merging all the independent peaks from gut and whole embryo ATAC-seq data resulting in 30866 open chromatin regions (OCRs) (Table S1). Majority of those OCRs are in the intergenic regions (40.44%) and in the introns (28.27%), promoter regions have 12.95% of the open chromatin peakset (Fig. 1A). We have also performed TF footprinting analysis to find which TFs could be bound at those OCRs. This resulted in 279197 footprints genome-wide (Table S2), with 36.06% of them in the intergenic OCRs, 27.10% in the intronic OCRs and 17.10% in the promoter-TSS associated OCRs (Fig. 1B). In order to assess whether the OCRs correspond to pCRMs, we have looked for the presence of published confirmed cis-regulatory regions in our consensus peakset. To do this, we have looked for overlap between the confirmed published hindgut CRMs (Arnone et al., 1998; Cui et al., 2017; de-Leon and Davidson, 2010; Lee et al., 2007; Livi and Davidson, 2007; McCarty and Coffman, 2013; Smith et al., 2008) and our 2 dpf stage consensus peaks. We have found that out of 22 confirmed published CRMs, 18 are present in the consensus peak set (Fig. 1C). For instance, our data recovers all the three known CRMs for *Sp-Blimp1*: regions CR3, CR5 and 43 (Smith et al., 2008) (Fig. 1D). Our data also improves resolution of CRM prediction since the open chromatin and thus active region of CRM E of *Sp-Hox11/13b* is confined to a 503 bp long peak inside of 4988 bp of the identified CRM E (Cui et al., 2017). This indicates that the consensus peakset represents pCRMs for the 2 dpf stage embryos. Some of the known CRMs were not found, such as CRM D for *Sp-Hox11/13b (Cui et al., 2017)* (Fig. 1D), which could be due to these locations being open at different stages of development and, therefore, were not further analyzed in this study. Our dataset also allowed us to identify novel CRMs as exemplified by pCRMs around the key posterior gut patterning drivers *Sp-Pdx1* and *Sp-Cdx* (Annunziata and Arnone, 2014; Cole et al., 2009). Specifically, there are five putative CRMs for *Sp-Pdx1* and four for *Sp-Cdx* at 2 dpf (Fig. 1E). *Sp-Pdx1* CRMs will be further explored in later sections of this manuscript.

**Figure 1.**
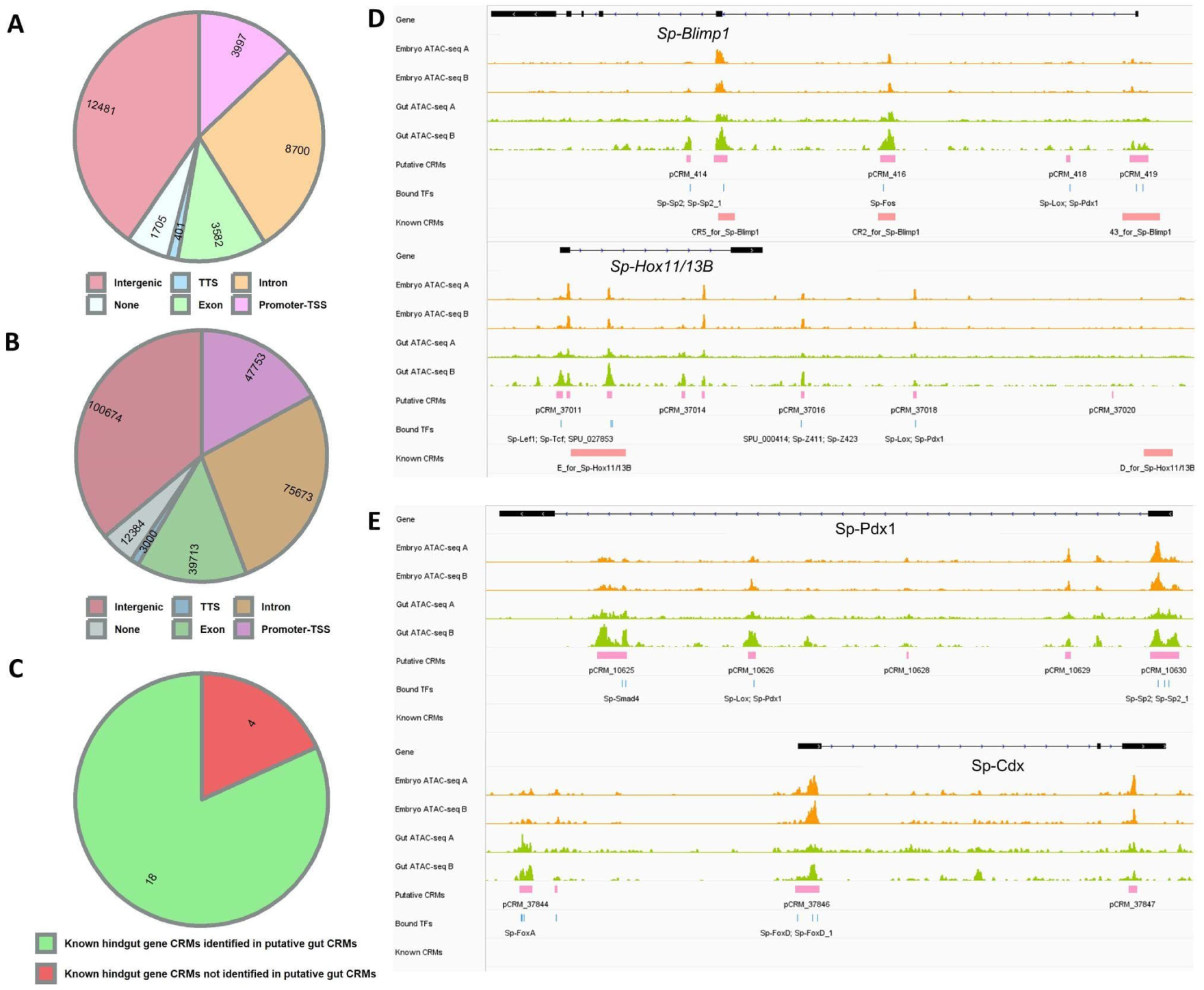
Open chromatin regions in the sea urchin gastrula. **A)** Pie chart of the proportion of putative CRMs relative to the genome annotation features. **B)** Pie chart of the proportion of transcription factor footprints in pCRMs associated with genome annotation features. **C)** Pie chart of the proportion of presence of published known hindgut CRMs in the ATAC-seq predicted putative CRMs at 2 dpf. **D)** Genome browser tracks of *Sp-Blimp1* and *Sp-Hox11/13b* putative CRMs overlapping with published CRMs for those genes. **E)** Genome browser tracks of *Sp-Pdx1* and *Sp-Cdx* putative CRMs. No CRMs were previously published for these genes.

### Diversity of cell type families in the sea urchin gastrula

In order to identify cell type specific GRNs in the 2 dpf *S. purpuratus* late gastrula stage we performed scRNA-seq on five samples originating from three independent biological replicates. Embryos were dissociated into single cells using an enzyme-free dissociation protocol, previously developed in our group (Paganos et al., 2021), which allows the sequencing of alive and healthy cells. Isolated single cells were processed using the 10x Chromium scRNA-seq capturing system (Fig. 2A). In total, transcriptomes from 15,341 cells were included in the final analysis. Computational analysis, including data integration and Louvain graph clustering resulted in the identification of 20 distinct cell clusters (Figs 2A and 2E), corresponding to individual cell types or a set of closely related cell types.

**Figure 2.**
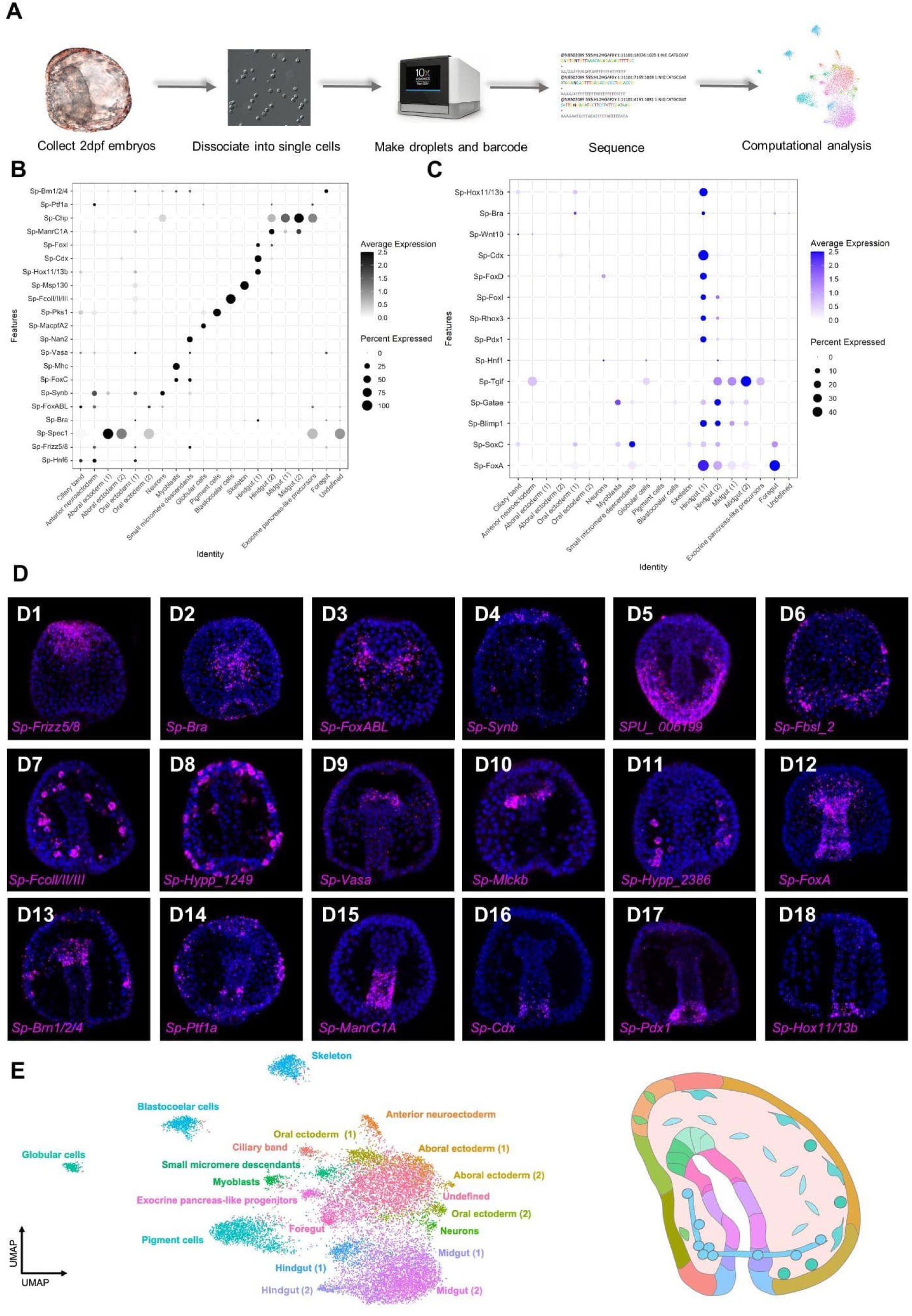
Cell type family atlas of the 2 dpf *S. purpuratus* gastrula stage. **A)** General scheme of scRNA-seq data generation and analysis. **B)** Dotplot of marker genes used for determining cluster identities. **C)** Dotplot of known hindgut genes, highlighting expression in the Hindgut (1) cluster. **D)** FISH of *S. purpuratus* 2 dpf gastrula embryos using antisense probes for *Sp-Frizz5/8* (D1), *Sp-Bra* (D2), *Sp-FoxABL* (D3), *Sp-Synb* (D4), *SPU_006199* (D5), *Sp-FcoI/II/III* (D6), *Sp-Hypp_1249* (D7), *Sp-Vasa* (D9), *Sp-Mlckb* (D10), *Sp-Hypp_2386* (D11), *Sp-FoxA* (D12), *Sp-Brn1/2/4* (D13), *Sp-Ptf1a* (D14), *Sp-ManrC1A* (D15), *Sp-Cdx* (D16), *Sp-Pdx1* (D17), *Sp-Hox11/13b* (D18). All embryos are oriented in oral view, except embryos shown in panels D7 and D14 that are oriented in lateral view and D5 that is placed in dorsal view. Nuclei are depicted in blue (DAPI). **E)** Uniform Manifold Approximation and Projection (UMAP) plot of identified cell clusters in 2 dpf gastrula, with the clusters highlighted in a schematic representation of 2 dpf sea urchin gastrula using color-coding.

Next, we undertook the task to explore the identity of the 20 identified cell clusters. Cell type family identities were assigned to each cluster based on the expression of previously described cell type markers, exploration of the total amount of genes expressed within them (Table S3), as well as taking advantage of the transcriptional signatures previously identified in the 3 dpf pluteus larva through scRNA-seq (Paganos et al., 2021) (Fig. 2B, Fig. S1). For instance, the following genes were used as specific cell type markers allowing us to recognize distinct cell type families: ciliary band (*Sp-Hnf6, Sp-Fbsl_2*) (Paganos et al., 2021; Poustka et al., 2004), anterior neuroectoderm (*Sp-Frizz5/8*) (Cui et al., 2014), aboral ectoderm (Spec1, SPU_006199) (Amore et al., 2003; Paganos et al., 2021), oral ectoderm (*Sp-Bra* and *Sp-FoxABL*) (Paganos et al., 2021; Wei et al., 2012), neurons (*Sp-Synb*) (Burke et al., 2006), myoblasts (*Sp-FoxC, Sp-Mlckb*) (Andrikou et al., 2013; Paganos et al., 2021), small micromere descendants (*Sp-Nan2* and *Sp-Vasa*) (Juliano et al., 2010), globular cells (*Sp-MacpfA2*) (Ho et al., 2017), immune cells (*Sp-Pks1, Sp-Hypp_1249)* (Paganos et al., 2021; Perillo et al., 2020), blastocoelar cells (*Sp-FcolI/II/III*) (Paganos et al., 2021), skeleton (*Sp-Msp130, Sp-Hypp_2386*) (Harkey et al., 1992; Paganos et al., 2021), hindgut (*Sp-Hox11/13b*, *Sp-Cdx*, *Sp-FoxI*) (Annunziata and Arnone, 2014; Tu et al., 2006), midgut (*Sp-Chp*, *Sp-ManrC1a*) (Annunziata and Arnone, 2014; Annunziata et al., 2019), exocrine pancreas-like precursors (*Sp-Ptf1a*) (Paganos et al., 2022b; Perillo et al., 2016) and foregut (*Sp-Brn1/2/4*) (Cole and Arnone, 2009). As the ultimate goal of this study is the reconstruction of the posterior gut GRN and especially the one that is orchestrated by the TF Sp-Pdx1, we further explored our data by plotting for *Sp-Pdx1* as well as TFs known to pattern distinct domains of the posterior archenteron (Annunziata and Arnone, 2014; Paganos et al., 2021) (Fig. 2C). Our analysis revealed that the majority of these TFs are expressed in the two clusters that we recognized as hindgut, supporting our cell type annotation. Interestingly, transcripts for *Sp-Pdx1* and, in total, eleven out of the thirteen investigated TFs were found present in the Hindgut (1) cluster. In more detail, the molecular fingerprint of this cell type family, as revealed by scRNA-seq, includes the TFs Sp-FoxA, Sp-SoxC, Sp-Blimp1, Sp-Gatae, Sp-Pdx1, Sp-Rhox3, Sp-FoxI, Sp-FoxD, Sp-Cdx, Sp-Bra and Sp-Hox11/13b. Furthermore, the presence of the TFs Sp-Bra and Sp-Hox11/13b in this cluster, known to be expressed in the most posterior part of the developing archenteron, confirms that this cell type family corresponds to the most posterior hindgut domain, even though *Sp-Pdx1* and *Sp-Bra* are not expressed in the same cells of this domain, as revealed by FISH (Fig. S2). This suggests heterogeneity of the Hindgut (1) cluster containing cells corresponding to more than one cell type.

Finally, to confirm the identities assigned to the cell clusters, we performed *in situ* hybridization on a selected set of genes with known and unknown expression patterns at this stage (Fig. 2D). The FISH results were in line with the initial predictions and our cell type annotation. Similar to our previous findings in the case of the 3 dpf *S. purpuratus* pluteus larva (Paganos et al., 2021), there is one cluster with poorly defined molecular signature that likely represents not fully differentiated cells, which we refer to as undefined. The persistence of this cluster at both timepoints is indicative of this cluster representing an actual cell type rather than it being a technical artifact.

### RNA-seq data uncovers the targets of Sp-Pdx1 in the developing gut

Causal information about the function of a gene is crucial to drafting GRNs. Taking advantage of the already available bulk RNA-seq data after Pdx1 knockdown at 2 dpf gastrula obtained from Annunziata and Arnone (2014), after re-analysis using up-to-date differential expression analysis pipeline, we were able to get the list of differentially expressed genes (DEGs) between the untreated sea urchin embryos and those that were injected with *Sp-Pdx1* morpholino oligonucleotides (MO) (Cole et al., 2009). The re-analysis of this dataset allowed us to provide additional information regarding these libraries. The principle component analysis (PCA) performed on the untreated and injected groups of embryos showed that 97% of variance between the samples is explained by the injection of the *Sp-Pdx1* morpholino (Fig. 3A), which indicates that the difference in gene expression is due to the *Sp-Pdx1* perturbation and, thus, the dataset can be used to infer causal information downstream of Sp-Pdx1. In line with previous findings, our differential expression analysis identified a large number of deregulated genes totaling to 2680 DEGs with adjusted *p*-value of less than 0.05 (Table S4). Of those, 1661 were upregulated after *Sp-Pdx1* perturbation and 1019 were downregulated (Fig. 3B). According to scRNA-seq, *Sp-Pdx1* positive cells are found in the hindgut (clusters Hindgut (1) and Hindgut (2)), in the midgut (clusters Midgut (1) and Midgut (2)) as well as a low number of cells in the clusters corresponding to oral ectoderm (oral ectoderm cells (2)) and ciliary band (Fig. 3D). Notably, the average scaled expression of this gene in clusters other than Hindgut (1) is less than 0.5 (Fig. 2C, Table S3). The cell type families of differentially expressed genes could be inferred using the scRNA-seq data (Fig. 3C). The downregulated genes, thus those that are activated by Sp-Pdx1 in the untreated embryos, belong to gut cell families, especially to the hindgut cell clusters, while the upregulated genes belong mostly to the anterior neuroectoderm and ciliary band cell clusters, although there are some genes in the anterior neuroectoderm that are downregulated after Sp-Pdx1 MO (Fig. 3C, E). Consequently, the RNA-seq data sheds light on what genes are downstream of Sp-Pdx1 and whether they are up or downregulated by this TF. This information can be used to iteratively predict causal information of the interactions suggested by the combination of ATAC-seq and scRNA-seq datasets.

**Figure 3.**
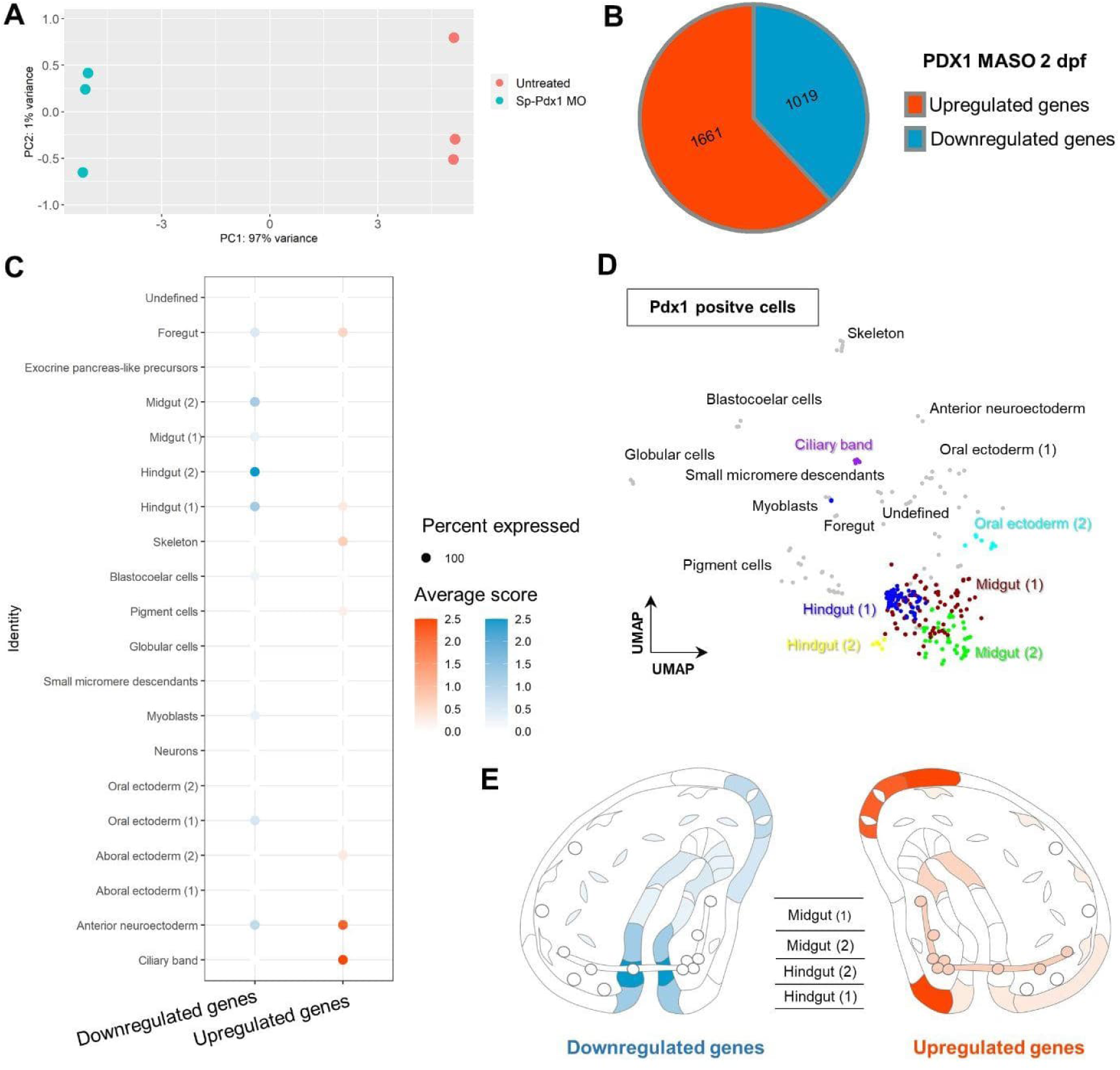
Differential RNA-seq of Sp-Pdx1 knockdown embryos. **A)** Principal component analysis (PCA) plot highlighting the variance explained by the morpholino treatment. **B)** Pie chart of proportion of significant genes up- and down-regulated after Sp-Pdx1 knockdown. **C)** Dotplot of cell clusters to which up- and down-regulated genes belong. **D)** UMAP plot of cells within the identified cell clusters at 2 dpf that express *Sp-Pdx1*. **E)** Diagrams of sea urchin 2 dpf gastrula highlighting the localization of up- and down-regulated genes.

### Gastrula hindgut Gene Regulatory Network

Our omics data allows drafting GRNs for any individual cell type in the 2 dpf *S. purpuratus* gastrula. For instance, the complete GRN for the 2 dpf gastrula hindgut can be drafted, in particular, the interactions between various TFs involved in building the embryonic hindgut with 512 interactions involving 91 individual TFs (Fig. S3A, Table S5). This GRN can be narrowed down keeping only the nodes that are co-expressed with *Sp-Pdx1* in the same cells (Fig. S3B, Table S6).

Previously, the core of the hindgut GRN was reconstructed by Annunziata and Arnone (2014) and a global genome view of the GRN operating within the nuclei of these cells was reviewed in Annunziata et al. (2019). Our combinatorial data allows us to update and refine the published GRN for the hindgut region and reconstruct the one for Sp-Pdx1 positive cells at higher resolution. The GRN presented in Annunziata et al. (2019) (Fig. 4A), which represents the most complete version of the posterior gut at that time, shows possible interactions between the different genes comprising it. However, with the available information at that point it was unclear whether these interactions could be direct or indirect. Using our datasets towards answering the very same question, we found that most of them appear to be indirect and each gene is wired through one or more intermediate nodes. Bellow we report several cases for which we present a refined and/or alternative gene regulatory connectivity:

**Figure 4.**
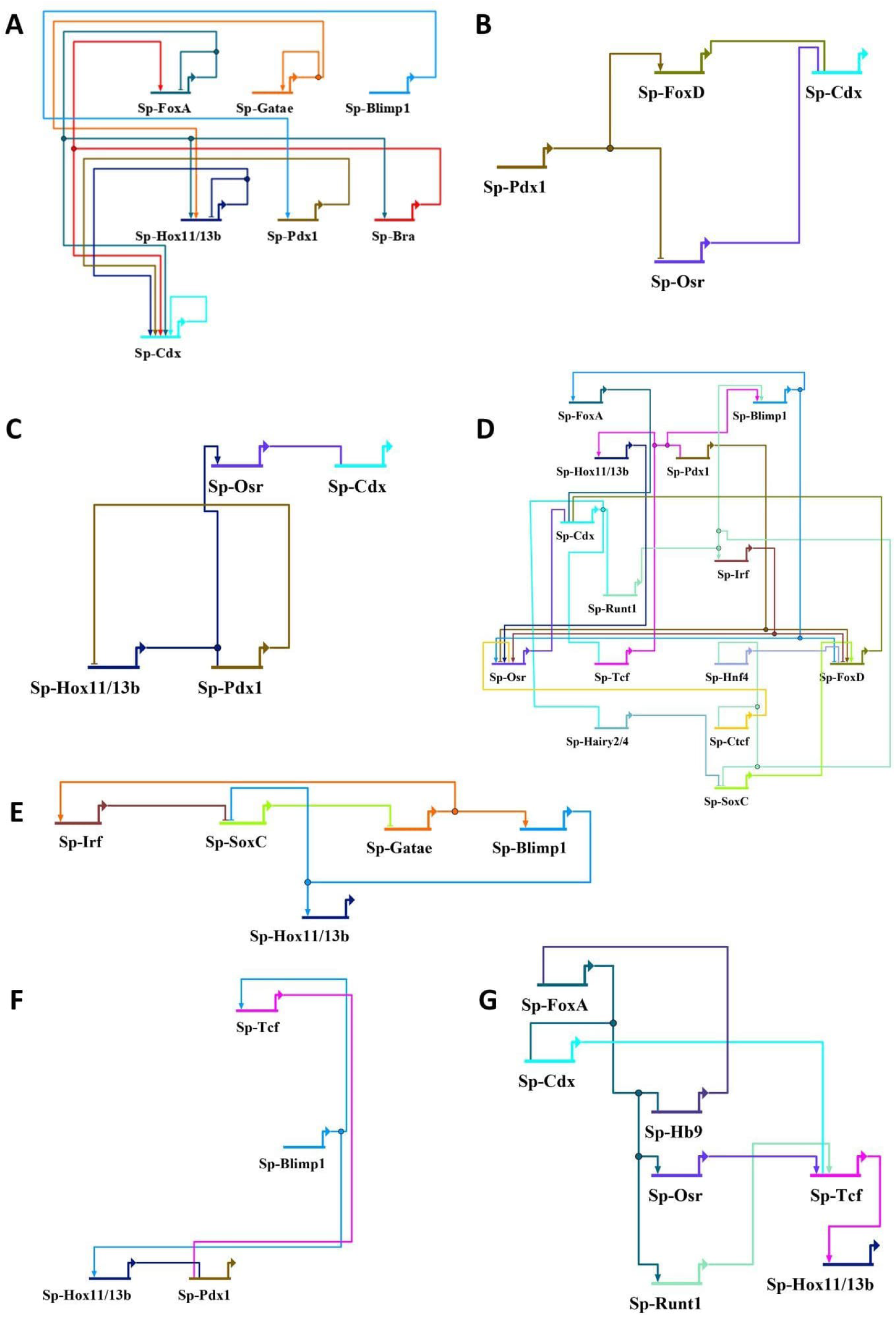
2 dpf *S. purpuratus* gastrula stage hindgut gene regulatory networks. **A)** Global genome view of the core TFs operating in the 2 dpf hindgut GRN as reviewed in Annunziata et al. (2019). **B)** *In silico* drafted GRN connecting Sp-Pdx1 with downstream genes of the core GRN. **C)** *In silico* drafted GRN connecting Sp-Hox11/13B with downstream genes of the core GRN. **D)** *In silico* drafted GRN connecting Sp-Cdx with downstream genes of the core GRN. **E)** *In silico* drafted GRN connecting Sp-Gatae with downstream genes of the core GRN. **F)** *In silico* drafted GRN connecting Sp-Blimp1 with downstream genes of the core GRN. **G)** *In silico* drafted GRN connecting Sp-FoxA with downstream genes of the core GRN.

#### Sp-Pdx1

Sp-Pdx1 was shown to control Sp-Cdx (Fig. 4A), however the actual link between the two remained an open question. Our data suggests that this interaction is likely indirect since there are no Sp-Pdx1 TF binding sites above the threshold near *Sp-Cdx*. This interaction may instead be routed through either Sp-FoxD or Sp-Osr (Fig. 4B) as revealed by this study.

#### Sp-Hox11/13b

Sp-Hox11/13b was previously shown to have a self repressive loop and an input on *Sp-Cdx* (Fig. 4A). The combinatorial data, again, favors the indirect nature of the self control loop. This loop can instead be wired through Sp-Pdx1, which represses *Sp-Hox11/13b* expression. Additionally, the effect Sp-Hox11/13b on *Sp-Cdx* can be explained through Sp-Hox11/13b direct activation of *Sp-Osr*, which then in turn affects *Sp-Cdx* directly. We also identify a direct input of Sp-Hox11/13b on *Sp-Pdx1*, which was not reported before (Fig. 4C).

#### Sp-Cdx

Similarly to *Sp-Hox11/13b*, *Sp-Cdx* also has a self regulatory loop (Fig. 4A). In this case our combinatorial analyses also suggest multiple equivalent indirect paths from Sp-Cdx back to *Sp-Cdx* that is in line with the *Sp-Cdx* self regulation previously reported, such as through Sp-Runt1 to Sp-Blimp1 onto *Sp-FoxD* and then to *Sp-Cdx* (Fig. 4D).

#### Sp-Gatae

Sp-Gatae was shown to have an activatory effect on *Sp-Hox11/13b* and on itself (Fig. 4A). Our data indicates that these effects are also indirect, and that the path to *Sp-Hox11/13b* can go through upregulating *Sp-Blimp1*, which itself upregulates *Sp-Hox11/13b*. The effect on *Sp-Hox11/13b* can also be explained through the same intermediates that are likely to account for the self feedback loop of Sp-Gatae such as Sp-Irf and Sp-SoxC (Fig. 4E).

#### Sp-Blimp1

Sp-Blimp1 was previously shown to affect *Sp-Pdx1* (Fig. 4A), and it is likely that this interaction occurs through a single intermediate, either Sp-Tcf or Sp-Hox11/13b or possibly both. Thus, this interaction between Sp-Blimp1 and *Sp-Pdx1* is also not direct (Fig. 4F).

#### Sp-FoxA

In the Annunziata et al (2019) GRN, *Sp-FoxA* has many outputs: Sp-FoxA, Sp-Cdx, Sp-Hox11/13b and Sp-Bra (Fig. 4A). The input on *Sp-Cdx* could be a direct one, as supported by TF *in silico* footprinting. The data for the rest of the interactions suggests that they are indirect and happen through intermediates. Sp-FoxA is likely to affect itself through a loop via Sp-Hb9, while the input on *Sp-Hox11/13b* can be explained via Sp-Osr or Sp-Runt1 to Sp-Tcf activations. Sp-Tcf, eventually, has a direct input on *Sp-Hox11/13b* (Fig. 4G).

#### Sp-Bra

*Sp-Bra* is not expressed in the hindgut cells expressing *Sp-Pdx1* at the 2 dpf gastrula stage (Fig. S2) but going back to include all the cells of the Hindgut (1) cluster its wiring can also be recovered (Fig. S4). Our data confirms the direct input of Sp-Bra on *Sp-FoxA* with an Sp-Bra footprint proximally to the *Sp-FoxA* gene (de-Leon and Davidson, 2010; Schwaiger et al., 2022), while the Sp-Bra effect on *Sp-Cdx* is likely indirect and could go either through Sp-FoxA or Sp-Osr as intermediates. In addition, our data at 2 dpf suggest that the inputs of Sp-FoxA on *Sp-Bra* and on itself are indirect, the effect on *Sp-Bra* could be wired through Sp-Tcf and Sp-Cdx, Sp-Runt1 and Sp-Osr intermediates, while self-feedback loop of Sp-FoxA is to be explained by a loop through Sp-Tcf and Sp-Bra (Fig. S4).

All together, the presented omics datasets not only allow correct reconstruction of a GRN without prior knowledge, but also empowers better resolution of the interactions identified previously by supplementing additional information on whether these interactions are direct or indirect and by identifying the intermediate nodes within cells of interest through combinatorial analyses.

### CRM5 allows control of *Sp-Pdx1* by Sp-Hox11/13b

The identified direct input of Sp-Hox11/13b on *Sp-Pdx1* was not reported previously and, therefore, is of great interest. Consequently, we focused our attention on exploring this input. Our CRM predictions and the TF footprinting indicates a binding site for Sp-Hox11/13b in the pCRM5 which is in the 5’ of the gene and overlaps the first exon of the *Sp-Pdx1* gene (Fig. 5A). To determine whether this pCRM is indeed an active cis-regulatory element, a GFP reporter construct with this element was microinjected into *S. purpuratus* zygotes. The microinjected embryos were allowed to develop until 2 dpf gastrula stage and then GFP expression driven by this pCRM was visualized. At 2 dpf, the CRM5-GFP-tag construct exhibits expression consistent with the known Pdx1 positive domains including the lateral neurons, (Fig. 5B, G) and the hindgut regions (Fig. 5C, G) as well as other regions of the gut such as the future stomach (Fig. 5D, G). At 3 dpf pluteus stage, this CRM is capable of driving GFP expression in the predominant Pdx1 expression domain in the larval digestive tract, specifically, in the pyloric sphincter (Fig. 5E). To determine whether Sp-Hox11/13b could affect *Sp-Pdx1* expression through CRM5, Sp-Hox11/13b morpholino oligonucleotide, previously shown to strongly downregulate Sp-Hox11/13b function (Arenas-Mena et al., 2006), was co-injected with the CRM5 GFP reporter construct.

**Figure 5.**
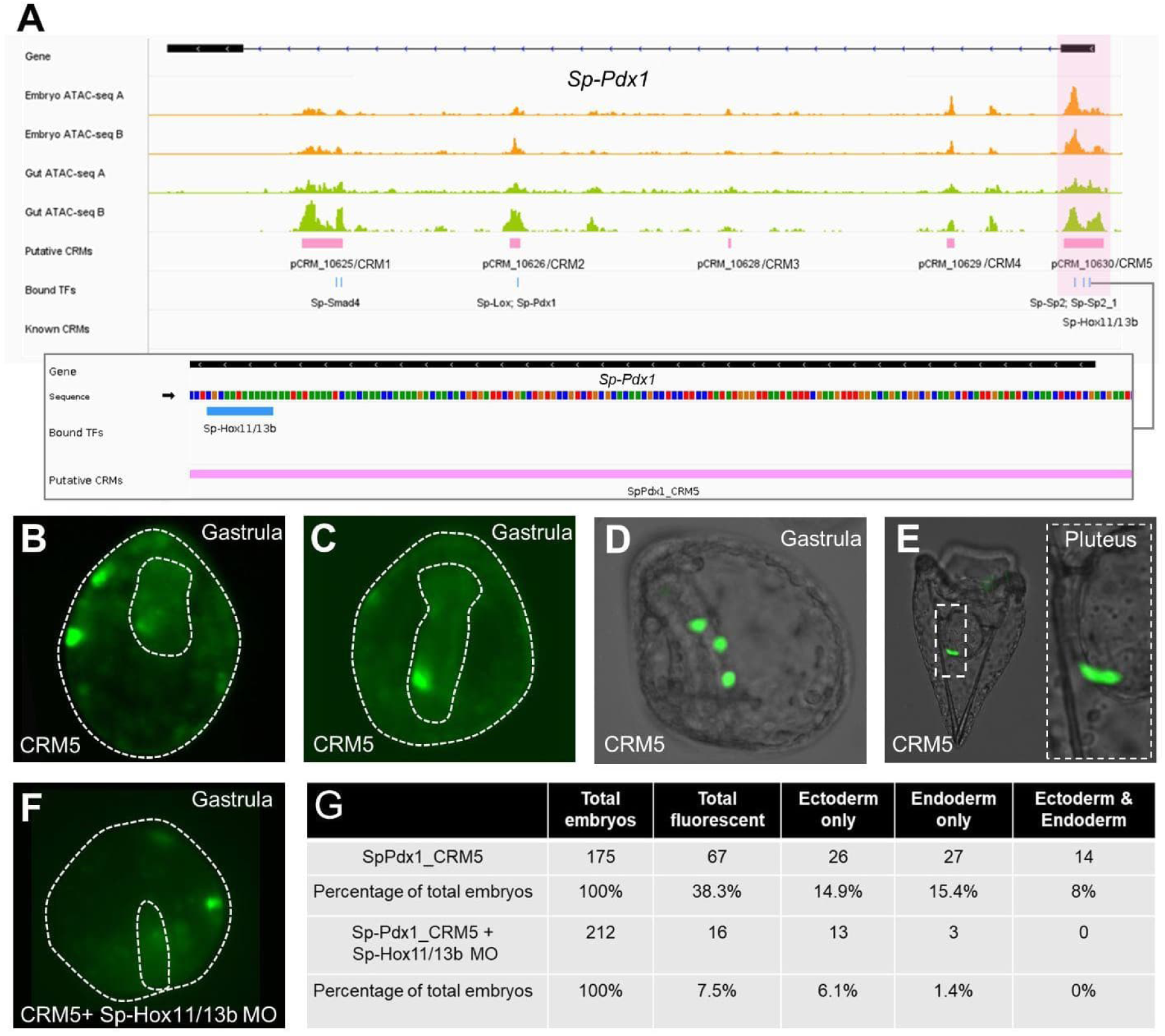
Sp-Pdx1 CRM5 control of *Sp-Pdx1*. **A)** Genome browser tracks of *Sp-Pdx1* putative CRMs, indicating position and sequence of Sp-Hox11/13b binding site within CRM5: the zoom-in shows the Sp-Hox11/13b binding site sequence using color-coding: A is green, T is red, C is blue and G is orange. **B)** Sp-Pdx1 CRM5 driven GFP expression in the lateral neurons of the 2 dpf gastrula stage. C) Sp-Pdx1 CRM5 driven GFP expression in the hindgut of the 2 dpf gastrula stage. D) Sp-Pdx1 CRM5 driven GFP expression in the midgut of the 2 dpf gastrula stage. **E)** Sp-Pdx1 CRM5 driven GFP expression in the midgut of the 3 dpf pluteus. **F)** Sp-Pdx1 CRM5 driven GFP expression in the 2 dpf gastrula stage indicating absence of endodermal expression after Sp-Hox11/13b knockdown. **G)** Embryo scoring table showing the effect of Sp-Hox11/13b knockdown on the Sp-Pdx1 CRM5 driven GFP expression in different germ layers.

At 2 dpf, the presence of the Sp-Hox11/13b morpholino decreases the overall reporter construct expression, in addition to greatly inhibiting the GFP expression in the endodermal regions, such as the hindgut (Fig. 5F, G), with more than ten-fold decrease in the percentage of endodermally expressing embryos (Fig. 5G). This indicates that Sp-Hox11/13b could control *Sp-Pdx1* expression in the hindgut through *Sp-Pdx1* CRM5.

### Multi-omics increases hindgut GRN resolution

Combinatorial omics and in vivo validations allow drafting an updated version of the GRN published in Annunziata and Arnone (2014) within the cells that express the Sp-Pdx1 transcription factor (Fig. 6). Chromatin accessibility information allowed us to identify putative CRMs and TF binding sites within them, while single cell data allowed us to filter those to contain only genes from cells that express the *Sp-Pdx1* gene. Such filtering to cells expressing a particular TF allows us to assess its role controlling downstream genes in the same cells, when adding the bulk RNA-seq data after *Sp-Pdx1* knockdown. This gives a highly detailed and novel GRN for a particular subset of hindgut cells expressing *Sp-Pdx1* cells. Notably, these cells express all the TFs present in the GRN reviewed in Annunziata et al. (2019) except *Sp-Bra*. The reasoning for this is that at 2 dpf *Sp-Bra* is expressed in a different subset of Hindgut (1) cells and is not co-expressed with *Sp-Pdx1*, thus it was excluded from the updated GRN (Figs 6, S2). *Sp-Hox11/13b* MO injected embryos showed a decrease in hindgut expression of a putative *Sp-Pdx1* CRM5-GFP construct that normally shows *Sp-Pdx1*-like expression pattern. This allowed us to validate the cis-regulatory direct effect of Sp-Hox11/13b on *Sp-Pdx1* via *Sp-Pdx1* CRM5 (Fig. 6 green diamond). The cis-regulatory effect of Sp-Tcf on *Sp-Hox11/13b* and *Sp-Blimp1* has been shown previously at blastula 18-24 hours post fertilization (Cui et al., 2017; Smith et al., 2008), and our data suggests that these effects persists in the 2 dpf gastrula as well (Fig. 6 orange diamonds). In addition, we have also shown that many of the previously identified interactions are indirect and happen via intermediates suggesting high complexity and depth of the 2 dpf sea urchin hindgut GRN even for nodes co-expressed with *Sp-Pdx1* (Fig. 6).

**Figure 6.**
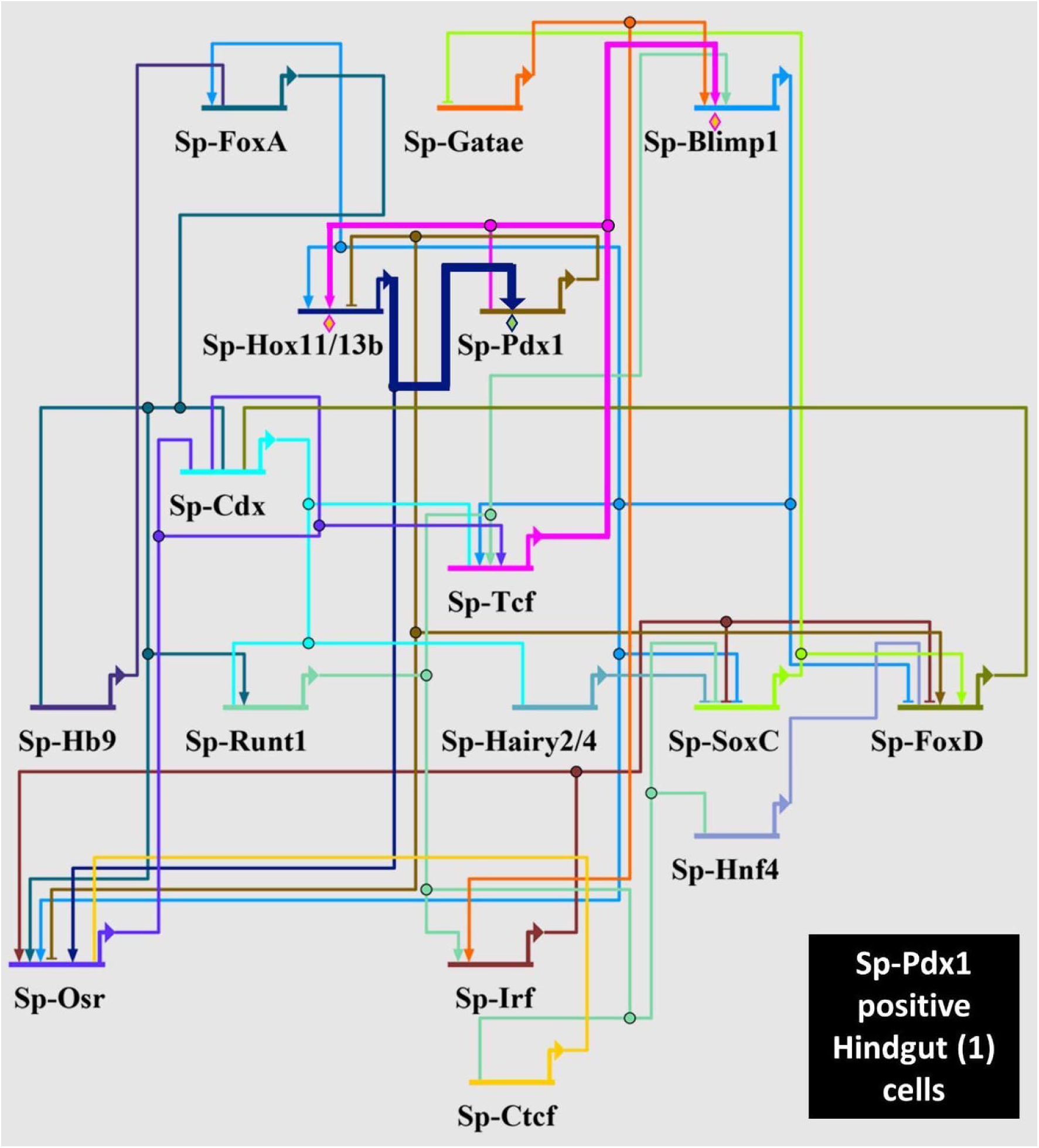
GRN operating in Sp-Pdx1 positive hindgut cells. Updated GRN of previously published nodes within Sp-Pdx1 positive hindgut cells drafted using the presented multi-omics approach. Validation of interactions by cis-regulatory analysis are indicated by orange diamonds (Cui et al., 2017; Smith et al., 2008).

## Discussion

### Predicted GRN recapitulates and refines existing GRN

The sea urchin endomesoderm GRN is one of the most studied. However, the GRNs were traditionally drafted for morphologically identifiable tissues which may or may not express all the nodes at a particular time point. In addition, historically, GRNs had limited information, mostly due to lack of high-throughput technologies at the time, on whether a particular interaction is direct or indirect, unless cis-regulatory analysis was performed. Lack of information as to the directness, allows only the overall final effect of perturbation on a gene to be identified. The integrative multi-omics allows to solve these issues. The GRN presented in this manuscript is the first GRN for a cell population expressing a particular TF, in this case *Sp-Pdx1*, and all the nodes are shown to be co-expressed with *Sp-Pdx1* in the same cells by scRNA-seq data. ATAC-seq data, on the other hand, points to which interactions of the GRN are likely to be direct and which are not, and through which intermediates they could go. These intermediates allow deducing the most parsimonious effect of a gene perturbation on the downstream targets, direct or indirect. Thus, the available omics data allows to tackle sea urchin GRNs at unprecedented resolution, which refines the existing GRNs and highlights the complexity and depth of the regulatory wiring responsible for the development of cell types, tissues and other embryonic structures. Our data suggests that the hindgut GRN, and likely, the whole endomesoderm GRN, is complex and diverse with many TF genes serving as intermediates in the interactions of the GRN nodes.

The published interactions of the hindgut GRN were recapitulated through our approach, albeit showing that most of them are likely indirect, increasing the resolution of the interactions of the nodes. This approach stays within the logic of Materna and Oliveri 2008 for drafting GRNs as the embryological knowledge is obtained via scRNA-seq data while the knowledge of the regulatory genes such as TFs as well as CRMs that establish the regulatory state are identified via ATAC-seq and scRNA-seq datasets. Finally, the causal information can be added through differential bulk RNA-seq after gene perturbation. Thus, all the necessary components for GRN drafting as per Materna and Oliveri 2008 can be obtained through multi-omics and their integration.

### Similarities and differences between sea urchin and vertebrate gut GRNs

The possibility of using vertebrate Position Weight Matrices (PWMs) for our TF footprinting also highlights the remarkable, albeit perhaps expected, conservation of the gut GRN nodes between sea urchin and vertebrates, which was previously noted (Annunziata et al., 2014; Annunziata et al., 2019; Grainger et al., 2010). However, despite the observed extreme conservation of topology of expression of TFs and signaling molecules, the exploration of the gene regulatory interactions connecting such factors so far mostly highlighted divergence in the architecture of the vertebrate and echinoderm gut GRNs (Arnone et al., 2016). One possible explanation of this apparent conundrum is that the GRN analyses were too shallow to reveal deeper homologies. The increased resolution of our refined sea urchin hindgut GRN allows now a deeper comparative analysis, reported below, of the genetic interactions between these homologous factors in these animals that evolutionarily diverged more than 570 MY ago (Yue et al., 2014).

The Sp-Pdx1 and Sp-Cdx are homologs of vertebrate PDX1 and CDX2, respectively, which have well-known functions in the vertebrate gut with both genes being crucial for gastro-intestinal tract development (Gao et al., 2009). While CDX2 controls intestinal development, PDX1 has a prominent role in the development of the vertebrate pancreas and duodenum. Analysis of PDX1 binding via ChIP-seq shows PDX1 binding sites near *CDX2,* which co-express in vertebrate duodenum *(Teo et al., 2015)*. ChIP-seq also suggests that PDX1 controls *GATA6* (potential homolog of Sp-Gatae), *HES1* (potential homolog of *Sp-Hairy2/4*), *FOXA2* (*Sp-FoxA*) and *MNX1* (*Sp-Hb9*) in human pancreatic progenitors or adult islet cells (Teo et al., 2015; Wang et al., 2018b). FOXA2 in turn can directly control PDX1 through its enhancers, suggesting a regulatory loop between the two genes (Gao et al., 2008). *HNF4*α (*Sp-Hnf4*) is also controlled by PDX1 in the human adult pancreas, which occurs through the P2 promoter (Thomas et al., 2001). These interactions appear in our updated GRN as well as Sp-Pdx1 has an effect on the sea urchin homologs of these genes, however most of them are indirect and happen through intermediates. *Sp-Cdx* could be controlled by Sp-Pdx1 via *Sp-FoxD*. *Sp-Gatae*, *Sp-Hairy2/4*, *Sp-Hnf4*, *Sp-FoxA* and *Sp-Hb9* are linked with Sp-Pdx1 through multiple intermediates such as *Sp-SoxC*, *Sp-Runt*, *Sp-Cdx*, *Sp-FoxD* and *Sp-Osr* (Fig. 6).

There is also data that RUNX3 (potential homolog of Sp-Runt1) protects mammalians from intestinal and gastric cancers by regulating *CDX2* and *PDX1* expression along with β-catenin/TCFs (Douchi et al., 2022; Ito, 2011), while the mammalian TCFs are potential homologs of Sp-Tcf. SOX4 and SOX12, which are potential homologs of Sp-SoxC, are co-expressed with PDX1 in endocrine pancreas cells, knockdown of these *SOX* genes does not seem to alter *PDX1* expression (Gracz and Magness, 2011; Wilson et al., 2005), but ChIP-seq revealed PDX1 bound near *SOX4 (Teo et al., 2015)* suggesting control of *SOX4* by PDX1. These interactions in our GRN also require intermediates and could again go through direct Sp-Pdx1 targets such as *Sp-Osr* and *Sp-FoxD* and then through *Sp-Cdx* to *Sp-Runt* or to *Sp-Hairy2/4* leading to *Sp-SoxC*. We predict that Sp-Runt has a direct activatory effect on *Sp-Tcf* and that together they can activate *Sp-Blimp1*. However, our data does not suggest the direct effect of Sp-Runt or Sp-Tcf on *Sp-Pdx1* or *Sp-Cdx* (Fig. 6).

CTCF plays the pivotal role in chromatin organization in vertebrate nuclei (Rowley et al., 2017) and was suggested to play a role in vertebrate GI tract development through regulating preproinsulin (*INS*) gene promoter organization which can lead to changes in binding of known *INS* promoter genes such as PDX1, HNF4α and NEUROD1 and this may happen for other GI tract CRMs (Babu et al., 2008; Wang et al., 2022; Xu et al., 2011). The roles of its sea urchin homolog Sp-Ctcf are not yet known. Knock-down experiments have suggested that sea urchin *H. pulcherrimus* CTCF orchestrates chromatin organization during mitosis of early development and its potential role in control of zygotic gene expression, but its exact role in gene regulation during later stages of embryogenesis is unclear, as knock-out of Hp-CTCF does not alter the development of the pluteus larva in *H. pulcherrimus* (Watanabe et al., 2023). Still, Sp-Ctcf might be implicated in the interactions we present in our updated GRN by organizing chromatin around CRMs of its target genes and allowing other TFs to bind these CRMs.

There are also regulatory relationships between *PDX1* and *OSR1 or OSR2* (Sp-Osr). *OSR1* is expressed in different regions of the gastro-intestinal tract such as foregut, stomach and pancreas (Han et al., 2020; Otani et al., 2014; Yu et al., 2019), while OSR2 is an intestinal marker (Zhao et al., 2022). OSR1 could affect *PDX1* indirectly in gastric tissues by repressing *TCF* transcription (Otani et al., 2014), while there are PDX1 ChIP-seq peaks near *OSR1* and *OSR2* genes (Teo et al., 2015). This could suggest, again, that OSR genes could control PDX1 and vice versa, thus paralleling the Sp-Pdx1 on *Sp-Osr* interaction in our updated GRN, which our data also suggests to be direct and not requiring intermediates. However, we did not detect Sp-Osr binding sites near *Sp-Pdx1*, therefore Sp-Osr on *Sp-Pdx1* interaction is also likely indirect (Fig. 6).

Gene to gene relationships between vertebrates PRDM1 (homolog of Sp-Blimp1), IRF1 (homolog of Sp-Irf), HOXD13 (Sp-Hox11/13b) and FOXD1 or FOXD2 or FOXD3 (Sp-FoxD) and PDX1 are unclear, which could be due to differences in expression localization as well as simply lack of research linking them on the molecular level. PRDM1 and IRF1 control the development of neonatal intestines, where PRDM1 acts in opposition to IRF1 to control immune response of the intestinal epithelium (Mould et al., 2015). BLIMP1 may also have a role in controlling pancreas development (Muraro et al., 2016) and is implicated in pancreatic cancers (Zhou et al., 2022). *HOXD13 (Kondo et al., 1996; Yahagi et al., 2004)*, *FOXD2 (Kim et al., 2023)* and *FOXD3 (Wang et al., 2018a)* are required for the development of the most posterior hindgut regions such as rectum and anus. Thus, expression of these genes in vertebrates may not co-localize with *PDX1*.

To summarize this analysis, it seems that for a lot of vertebrate homologs of the genes present in the updated hindgut GRN, PDX1 can exert direct control over via binding in the proximity to those genes (see above) in pancreas and the posterior regions of the gut. While the resolution of our updated gut GRN revealed complex interactions further highlighting the high degree of conservation of players previously observed (Pdx1, Cdx, FoxA, Blimp1 etc) and, in addition, suggested the involvement of TFs such as Osr, Runt, Hnf4, Irf etc, which were absent in the previous sea urchin gut GRN reconstruction, our data confirms that, overall, major rewirings occurred in the posterior gut GRN around Pdx1 between vertebrates and echinoderms.

### Limitations and strengths of the approach

The multi-omics approach used in this work has its limitations. Majority of TF position weight matrices are available for human or mouse TFs (Fornes et al., 2020) only with very few sea urchin PWM published. Thus, human homologs are used for TF footprinting, which could lead to false negatives or positives if the sea urchin homologs have different binding sequences. In addition, using vertebrate PWMs, certain TFs have similar PWMs such as forkhead genes (FoxD, FoxA etc) or SRY genes (e.g. SoxC), which complicates identification of TFs binding to a particular locus in a CRM and could also lead to incorrect identification of the homolog affecting a particular gene in other organisms such as vertebrates discussed in the previous section.

The tools available are designed for cell cultures, so data from whole tissue/embryo is likely to involve a lot of noise, and some gut interactions could potentially be hidden by reads coming from mesodermal cells still attached at the tip of the archenteron (presumptive coelomic pouches or muscles) that have a different TF repertoire bound to their DNA that could be left over during the gut tissue preparation. This could lead to false positives and false negatives. Thus, the indicative nature of our approach highlights the need for *in vivo* validations. With published CRM analysis data lacking for 2 dpf, experimental *in vivo* validations are necessary to validate the results of our approach at this stage of embryo development.

It is also important to note that our approach does not take into account cases in which transcription factors operate within a cell type, while they are not actively transcribed by its cells. Such are the cases of protein remainants from a previous developmental timepoint that could still be active, due to the longer protein half-life. For instance, *Sp-Pdx1* could affect more cells than the ones actively expressing it at 2 dpf gastrula stage as Sp-Pdx1 protein could persist in the cells and continue to function after active transcription and translation of this TF have ended. Thus, more cells of the hindgut could have genes under Sp-Pdx1 control if these cells were expressing *Sp-Pdx1* earlier in development. This could also explain the widespread, in terms of number of downregulated genes, effect of Sp-Pdx1 knockdown observed in cells of the Hindgut (2) cluster, as shown in Fig. 3C,D.

Despite the highlighted limitations, the importance of multimodal approaches to improve GRN inference has been recently highlighted (van der Sande et al., 2023). For vertebrate experimental systems, approaches using scRNA-seq, or combination of scRNA-seq and scATAC-seq of the same cells emerge (Yang et al., 2023; Zhang et al., 2023), which, similarly to our approach, allows drafting of single cell GRNs. These approaches are robust in identifying nodes and interactions of the GRNs, although they lack information on whether a given identified interaction is activatory or inhibitory. In our approach, we attempt to obtain the most parsimonious prediction to the directness of interactions predicted by ATAC-seq and scRNA-seq datasets using RNA-seq data. While the method presented in the current manuscript lacks single cell ATAC-seq data, the permissive nature of chromatin accessibility rather than deterministic allows us to use tissue level ATAC-seq data for GRN drafting, despite the potentially higher noise. Applying the omics data and integrating them as we present in the current manuscript gives the necessary information for GRN drafting at cell type level for the 2 dpf sea urchin embryo, while *in vivo* reporter construct experiments validate *in silico* predictions.

### Conclusions

Overall, our multi-omics approach drafts reliable GRNs that allow us to increase the resolution and highlight the high interconnectivity of the sea urchin hindgut GRN around *Sp-Pdx1*. Thus, within echinoderm research, such an approach has the potential of giving a more holistic view of gene regulation in a specific cell type in a developing embryo. Moreover, this approach is applicable to any experimental system, including emerging model organisms, as multi-omics datasets become more and more available thanks to the recent advances in NGS technologies.

## Materials and Methods

### Animal husbandry and culture of embryos

Adult *Strongylocentrotus purpuratus* individuals were obtained from Patrick Leahy (Kerckhoff Marine Laboratory, California Institute of Technology, Pasadena, CA, USA) and maintained in circulating seawater aquaria at Stazione Zoologica Anton Dohrn in Naples. Gametes were obtained by vigorously shaking the animals. Embryos and larvae were cultured at 15 °C in filtered Mediterranean Sea water diluted 9:1 with deionized water and collected 2 days post fertilization (dpf).

### Gut tissue separation

Sea urchin gut tissue was obtained by adapting existing protocols (Juliano et al., 2014; McClay, 2004). Echinoderm embryos were grown until the 2 dpf gastrula stage. The embryo suspensions were collected into 1.5 mL tubes using a 40 μm Nitex mesh filter and concentrated by spinning at 500 g for 5 min at 4°C and then washed once in 1 mL of Ca^2+^-Mg^2+^ free sea water (CMFSW). Then, the embryos were concentrated again by centrifuging at 500 g for 5 min at 4°C and treated with 1M glycine 0.02M EDTA in CMFSW for 10 minutes on ice. Embryos are pipetted up and down carefully using P1000 micropipette three times and transferred onto 1% agarose plates under the dissecting microscope to control the dissociation process. The guts were then collected individually into 1.5 ml Eppendorf tubes with 100 μl of artificial sea water on ice.

### ATAC-seq library synthesis

ATAC-seq libraries were generated as described in Magri et al. (2021). Around 270 gastrula stage embryos or 400 embryonic guts were collected per biological replicate and washed on the 40 μm Nitex mesh filter with artificial sea water (28.3g NaCl, 0.77g KCl, 5.41g MgCl2·6H2O, 3.42g MgSO4 or 7.13g MgSO4·7H2O, 0.2g NaHCO3, 1.56g CaCl2·2H2O per 1 L of deionized water) and transferred into 1.5 ml tubes. The samples were then centrifuged at 500 g for 5 min at 4 °C, washed twice with 200 μl artificial sea water in the 1.5 ml tube and then resuspended in 50 μl of lysis buffer (10 mM Tris-HCl, pH 7.4, 10 mM NaCl, 3 mM MgCl2, 0.2% IGEPAL CA-630), followed by lysis by pipetting up and down for 3-5 minutes. Half of the lysate was used for counting the released nuclei under Zeiss AxioImager M1 microscope using a haemocytometer with DAPI dye (1μl of 1:100 diluted DAPI in the 25 μl of the released nuclei). The other half was used for the tagmentation reaction by centrifuging the sample at 500 g, removing lysis buffer and then incubating for 30 minutes at 37°C with 25 μl of 2x tagmentation buffer (20 mM Tris-HCl, 10 mM MgCl2, 20% (vol/vol) dimethylformamide), 23.75 μl of nuclease free water and 1.25 μl of Tn5 enzyme. The Tn5 enzyme used for ATAC-seq library preparations was provided by Dr Jose Luis Gómez-Skarmeta.

The tagmented DNA was purified using MinElute Kit (Qiagen) following manufacturer’s instructions and eluted in 10μl of elution buffer. The eluted DNA was then amplified to obtain the library for sequencing (10 μl of eluted tagmented DNA, 10 μl of nuclease free water, 2.5 μl 10 μM Nextera Primer 1, 2.5 μl 10 μM Nextera Primer 2.X, where X is the unique Nextera barcode used for sequencing and 25 μl of NEBNext High-Fidelity 2x PCR Master Mix (New England BioLabs) using the following thermocycler program: 72°C for 5 min, 98°C for 30 sec, then 15 cycles of 98 °C for 10 sec, 63 °C for 30 sec and 72 °C for 1 min, followed by a hold step at 4°C. The amplified library was then purified with MinElute Kit (Qiagen) following manufacturer’s instructions and eluted in 20 μl of elution buffer. The concentration of the resulting library was checked using Qubit dsDNA BR Assay Kit (Molecular Probes) and its quality was assessed by running 70 μg of the library on 2% agarose 1x TAE gel. Libraries exhibiting nucleosomal size patterns on the gel were sent for sequencing. Sequencing was performed by BGI Tech (HongKong) resulting in 50 bp paired-end reads (Illumina HiSeq 4000) with the mean of 57 M reads.

### ATAC-seq data analysis

The raw ATAC-seq reads were trimmed from adapter sequences and bad quality bases using Trimmomatic 0.38 (Bolger et al., 2014) using the following parameters ILLUMINACLIP:TruSeq3-PE-2.fa:2:30:10:6 CROP:40 SLIDINGWINDOW:3:25 MINLEN:25 for paired end data and -phred33 quality scores. The good quality reads from each replicate are then mapped to the *S. purpuratus* genome v3.1 (Sea Urchin Genome Sequencing Consortium et al., 2006) using bowtie2 2.3.4.1 (Langmead and Salzberg, 2012). The mapped reads in BAM format were then converted into bed files using bedtools 2.27.1 (Quinlan and Hall, 2010), the bam file was filtered using samtools 1.7.1 (Li et al., 2009) to keep aligned reads of quality greater than 30, fix mates and remove duplicates with default parameters. In addition, only fragments of less than 130 bp in size were kept. The resulting files were then used in MACS2 2.1.2 software (Zhang et al., 2008) to call peaks using BED as input file format as well as setting --extsize to 100, --shift to 50 and using --nomodel setting with genome size of 815936258 for *S. purpuratus* genome v3.1; all the other MACS2 settings were kept at default. Bedtools 2.26.0 intersect tool (Quinlan and Hall, 2010) was used with default settings to combine replicates and obtain a consensus set of peaks, which were then all merged using bedtools merge to putative cis-regulatory modules (pCRMs) list.

The resulting gut ATAC-seq bam files were merged using bamtools and used as input in TOBIAS 0.11.1-dev (Bentsen et al., 2020) along with the genome and CRM list to perform transcription factor (TF) footprinting analysis. TOBIAS commands ATACorrect and ScoreBigwig were run with default parameters, BINDetect was run using JASPAR2020 vertebrate motif database position weight matrices (PWMs) (Fornes et al., 2020) using --motif-pvalue of 1e-5. Motifs for each TF with score of at least 0.5 were kept, to obtain a table with each pCRM and TFs bound to it. HOMER 4.10.3 (Heinz et al., 2010) annotatePeaks.pl was used to assign pCRMs to *S. purpuratus* genes. BLAST 2.6.0+ (Altschul et al., 1990) with max_target_seqs of 1 and max_hsps of 1 was used to identify single best sea urchin homolog for each of vertebrate TFs.

### Single cell embryo dissociation

Dissociation of the 2 dpf *Strongylocentrotus purpuratus* gastrulae into single cells was performed the same as in Paganos et al. (2021). Embryos were collected, concentrated using a 40 μm Nitex mesh filter and spun down at 500 g for 10 min. Sea water was removed and larvae were resuspended in Ca^2+^Mg^2+^-free artificial sea water. Embryos were concentrated at 500 g for 10 min at 4 °C and resuspended in a dissociation buffer containing 1 M glycine and 0.02 M EDTA in Ca^2+^Mg^2+^-free artificial sea water. Embryos were incubated on ice for 10 min and mixed gently approximately every 2 min, monitoring the progress of the dissociation. Dissociated cells were concentrated at the bottom of the tube at 700 g for 5 min and washed several times with Ca^2+^Mg^2+^-free artificial sea water. Cell viability was assessed using propidium iodide and fluorescein diacetate, samples with cell viability ≥ 90 % were further processed. Single cells were counted using a hemocytometer and diluted according to the manufacturer’s protocol (10X Genomics).

### Single-cell RNA sequencing and data analysis

ScRNA-seq sequencing and the analysis of the 2 dpf *Strongylocentrotus purpuratus* gastrula stage data was performed the same as in Paganos et al. (2021). Single cell RNA sequencing was performed using the 10X Genomics single-cell capturing system. Cells from three biological replicates, ranging from 6000 to 20,000 cells, were loaded on the 10X Genomics Chromium Controller. Single cell cDNA libraries were prepared using the Chromium Single Cell 3’ Reagent Kit (v3 and v3.1). Libraries were sequenced by GeneCore (EMBL, Heidelberg, Germany) for 75 bp paired-end reads (Illumina NextSeq 500). Cell Ranger Software Suite 3.0.2 (10x Genomics) was used for the alignment of the single-cell RNA-seq output reads and generation of feature, barcode and matrices. The genomic index was made in Cell Ranger using the *S. purpuratus* genome version 3.1 (Kudtarkar and Cameron, 2017; Sea Urchin Genome Sequencing Consortium et al., 2006). Cell Ranger output matrices for three biological and two technical replicates were used for further analysis in Seurat v3.0.2 R package (Stuart et al., 2019). Genes that are transcribed in less than three cells and cells that have less than a minimum of 200 transcribed genes were excluded from the analysis. The cutoff number of transcribed genes was determined based on feature scatter plots and varies depending on the replicate. In total 15,341 cells passed the quality checks and were further analyzed. Datasets were normalized and variable genes were found using the vst method with a maximum of 2000 variable features. Data integration was performed via identification of anchors between the five different objects. Next the datasets were scaled and principal component (PCA) analysis was performed. Nearest Neighbor (SNN) graph was computed with 20 dimensions (resolution 1.0) to identify the clusters. Transcripts of all genes per cell type were identified by converting a Seurat DotPlot with all these transcripts as features into a table (ggplot2 3.2.0 R package). All resulting tables containing the genes transcribed within different cell type families were further annotated adding PFAM terms (Finn et al., 2014; Trapnell et al., 2010) for associated proteins, gene ontology terms and descriptions from Echinobase (Kudtarkar and Cameron, 2017).

### Morpholino injections

Microinjections were performed as follows. The sea urchin eggs were dejellied in acidic seawater (pH 4.5) for <1 min, then washed in filtered sea water and rowed onto 4% protamine sulfate plates filled with PABA-FSW (50 mg of p-aminobenzoic acid in 100 ml of filtered sea water). The microinjecting needle was pulled from borosilicate glass with capillary using Sutter Instrument Co. Novato, CA using P-97 micropipette puller (Sutter), was loaded with the injection solution using a Microloader pipette tip (Eppendorf). Loaded needle tip was broken off using a scratch in the middle of the protamine plate with eggs. The eggs were fertilized with a few drops of diluted sea urchin sperm and injected with approximately 2 pL of microinjection solution. Injected eggs were washed twice with filtered sea water and incubated at 15 °C overnight, then the hatched embryos were transferred to 4-well plates (ThermoFisher Scientific) with filtered sea water to grow until 2 days post fertilization gastrula stage.

The morpholino oligonucleotide (MO) microinjection solutions containing contained 100 μM of final morpholino concentration were heated up to 75°C for 5 min and then passed through a 0.22 μm PVDF micro-filter (Millipore) placed in 500 μl tube by centrifugation at 2500 g for 2 min. After filtration, the filtrate was centrifuged for 15 min at top speed prior to microinjection. The translation morpholino sequence for *Sp-Hox11/13b* is the same as previously described (Arenas-Mena et al., 2006).

### RNA-seq data analysis

Raw reads for 2 dpf *Sp-Pdx1* MO injected and untreated embryos were published in Annunziata and Arnone (2014). The reads were re-analyzed for this manuscript, adhering to more up to date pipelines. Fastqc 0.11.5 (Andrews, 2010) was used to assess the quality of sequencing data. Bad quality sequences were trimmed from the reads using Trimmomatic 0.38 (Bolger et al., 2014) with the following options ILLUMINACLIP:TruSeq3-PE-2.fa:2:30:10:6 CROP:90 HEADCROP:10 SLIDINGWINDOW:3:25 MINLEN:25 using -phred33 base quality encoding for paired end reads. Only the paired output of Trimmomatic was used as input in Salmon 0.11.3 (Patro et al., 2017) for transcript quantification with automatic library detection and all settings left as default. Salmon index was made using transcriptome FASTA file corresponding with *S. purpuratus* genome version 3.1 from Echinobase.org (Cary et al., 2018; Tu et al., 2012) with k value of 25. The Salmon output quants files were used in DESeq2 (Love et al., 2014) R package for differential gene expression analysis; tximport (Soneson et al., 2015) package was used to link every transcript to a WHL ID denoting genes in the genome annotation. The analysis was performed using local fit and ∼condition (untreated or MO injected) as the factor. Differentially expressed transcripts with p-adjusted value of less than 0.05 using independent hypothesis weighting (IHW) were considered significant. Resulting transcripts were annotated adding *S. purpuratus* gene names.

### *In silico* GRN drafting

The general flowchart of the method is described in Fig. S5. The table with each putative sea urchin CRM and TFs bound to it, along with their target genes (described in ATAC-seq data analysis) was filtered to have only TFs and target genes present in a single cell type family. For this, the tables containing the genes transcribed within different cell type families, obtained in scRNA-seq data analysis as described above, were cut to have only genes expressed in the hindgut cluster. This list was used for filtering of the ATAC-seq analysis results. The resulting table of TFs and target genes was further filtered to have only TFs (Lambert et al., 2018) both as target and effector. In order to build a network downstream of Sp-Pdx, the data was further narrowed downto contain only genes expressed in Sp-Pdx positive cells. For this, AddModuleScore from Seurat (Stuart et al., 2019) was used to assess expression of Sp-Pdx1 in the 2 dpf cells; only cells where the resulting value was greater than 1.5 belonging to the hindgut cells cluster were taken for further analysis. Genes expressed in those cells with scaled average expression greater than 0.5 were used to build the GRN downstream of Sp-Pdx1 positive cells. To predict causal information of the interactions within the Sp-Pdx1 network results of the bulk RNA-seq differential expression analyses between untreated embryos and Sp-Pdx1 morpholino injected embryos were used. With the help of a custom R 4.1.2 (R Core Team, 2021) script direct targets of Sp-Pdx1 (based on ATAC-seq data) within Sp-Pdx1 positive cells were assigned causal dynamics information, then the targets of the direct targets, then the targets of indirect targets. This process was performed iteratively to give most parsimonious causal information to the genes downstream of Sp-Pdx1 in the GRN through iterative assignment of interaction signs. For instance, the Sp-Pdx1 gene in this case is the effector gene (Efct) that has direct (D-tgt) and indirect (I-tgt) targets (Fig. S6). If a direct target of Sp-Pdx1 (D-tgt1) is expressed less when Sp-Pdx1 is knocked down and its direct target, thus indirect Sp-Pdx1 target (I-tgt1) is also expressed less when Sp-Pdx1 is knocked down, then Sp-Pdx1 TF activates the D-tgt1 and D-tgt1 in turn activates I-tgt1 (Fig. S6). While, if an indirect Sp-Pdx1 target (I-tgt2), which is a direct target of a direct Sp-Pdx1 target, is downregulated by Sp-Pdx1 MO, thus upregulated by Sp-Pdx1 TF, and the direct Sp-Pdx1 target (D-tgt2) is downregulated by Sp-Pdx1 TF, then we can deduce that D-tgt2 downregulates the I-tgt2 (Fig. S6). Same approach can be used with the combination of up-regulating and down-regulating interactions, such as activation of I-tgt3 by Sp-Pdx1 (Fig. S6). This process can be performed iteratively for other indirect targets that have more intermediates between them and Sp-Pdx1 to assign signs to interactions downstream of Sp-Pdx1. The IDs used throughout the pipeline are then converted to gene names for visualization in BioTapestry software.

### CRM GFP reporter construct synthesis

Tagging of putative CRMs was performed according to a protocol described by Nam et al. (2010). The plasmids with DNA tags were provided by Dr Jongmin Nam. 50 bp flanks from the genome were added to the pCRMs to design primers using Primer3web 4.1.0 (Untergasser et al., 2012) falling within the added 50 bp to ensure the whole CRM is amplified. 18bp of the reverse complement of the beginning of the DNA tag sequences was added to the 5’ of the reverse primer so that the CRM can be combined with the DNA tag in equal amount using overlap PCR (Xiong et al., 2006) using Expand High Fidelity PLUS PCR (Sigma). The resulting fragment was run on 2% agarose 1x TAE gel, the corresponding band was cut out, and gel purified using GenElute Gel Extraction Kit (Sigma) according to manufacturer’s guidelines and eluted in 50 μl of kit elution buffer. The eluted DNA was then purified again using QIAquick PCR Purification Kit (Qiagen) and eluted in 30 μl of kit elution buffer. DNA yield and concentration was assessed using NanoDrop ND-1000 (ThermoFisher Scientific).

### CRM GFP reporter construct microinjections

The tagged CRM was used to make microinjection solutions according to Nam et al. (2010): 0.5 μl of tagged CRM solution, 1.2 μl of 1 mM KCl, 0.275 μl carrier DNA (genomic DNA sheared with HindIII enzyme (2 units for μg of DNA, in SuRE/Cut Buffer B for 3 h at 37 °C), purified using QIAquick PCR Purification Kit (Qiagen) and diluted to 500 ng/μl) and water up to 10 μl (Arnone et al., 2004; Nam et al., 2010). The prepared solutions were then centrifuged at top speed for 15 min until microinjections. The microinjection procedure was the same as described in Morpholino injections.

### Fluorescent *in situ* hybridization (FISH)

FISH was performed as described in Paganos et al. (2022a). In brief, embryos were collected 2 dpf and fixed in 4% PFA in MOPS Buffer overnight at 4°C. The following day specimens were washed with MOPS buffer several times and then stored in 70% ethanol at −20°C. Antisense mRNA probes were generated as described in Perillo et al. (2021). The primer sequences used for cDNA isolation were the same as previously described: *Sp-Synb (Burke et al., 2006)*, *Sp-FoxA*, *Sp-Pdx1, Sp-Brn1/2/4, Sp-Cdx, Sp-ManrC1a, Sp-Hox11/13b* (Annunziata and Arnone, 2014); *Sp-FoxC* (Andrikou et al., 2015); *Sp-Ptf1a* (Perillo et al., 2016); *Sp-Pks1 (Perillo et al., 2020)*; SPU_006199, Sp-Fbsl_2, S*p-Bra, Sp-FoxABL, Sp-Frizz5/8* and *Sp-FcoIl/II/III, Sp-Hypp_1249, Sp-Mlckb, Sp-Hypp_2386* (Paganos et al., 2021). Probes were synthesized from linearized DNA and labeled during synthesis with digoxigenin-11-UTP (Roche) nucleotides. Fluorescent signal was developed using the fluorophore conjugated tyramide technology. Specimens were imaged using a Zeiss LSM 700 confocal microscope.

## Acknowledgements

We thank Elijah Kareem Lowe for setting the foundation for designing the approach pipeline and Lorena Buono for help with scripts. We thank Rossella Annunziata for invaluable feedback on the manuscript and Davide Caramiello for taking care of the animals. We also thank Detlev Arendt, Jacob M. Musser and EMBL GeneCore for single cell transcriptomics.

## Data availability

The raw ATAC-seq and scRNA-seq sequencing data generated for this work and described in this manuscript can be found at NCBI Gene Expression Omnibus (accession numbers TBD). The bulk RNA-seq data from Annunziata and Arnone (2014) used in this manuscript can be found at https://osf.io/cbsxr/files/. Scripts associated with the data analysis pipeline can be found at https://github.com/Danvor/spur_2dpf_hindgut_pdx1_downstream_grn.

## Supplementary Figures

**Supplementary Figure 1.**
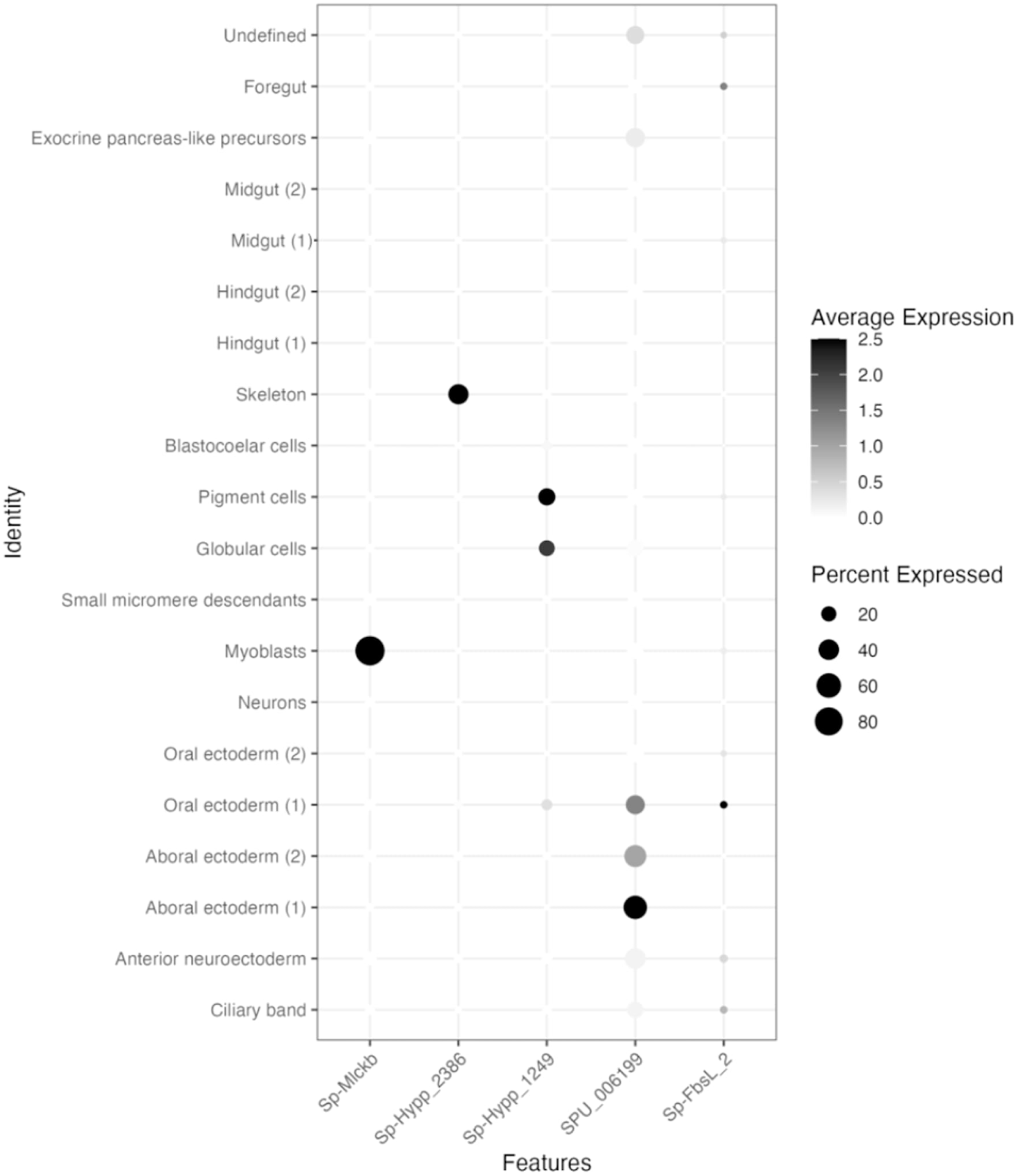
Dotplot of additional marker genes used for determining cluster identities.

**Supplementary Figure 2.**
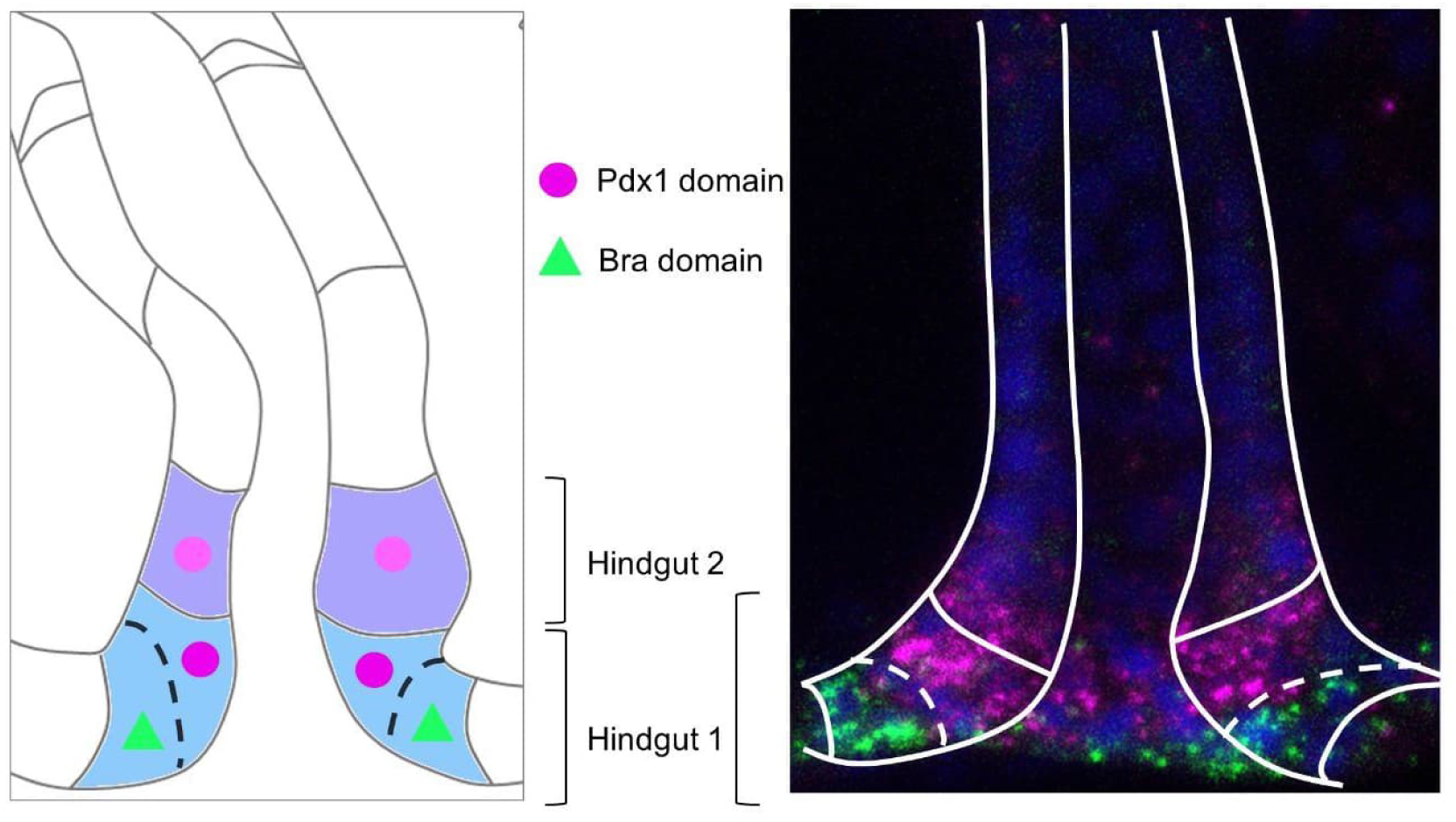
Schematic representation (left panel) and double FISH (right panel) showing *Sp-Pdx1* and *Sp-Bra* expression in different cells of the Hindgut (1) cluster.

**Supplementary Figure 3.**
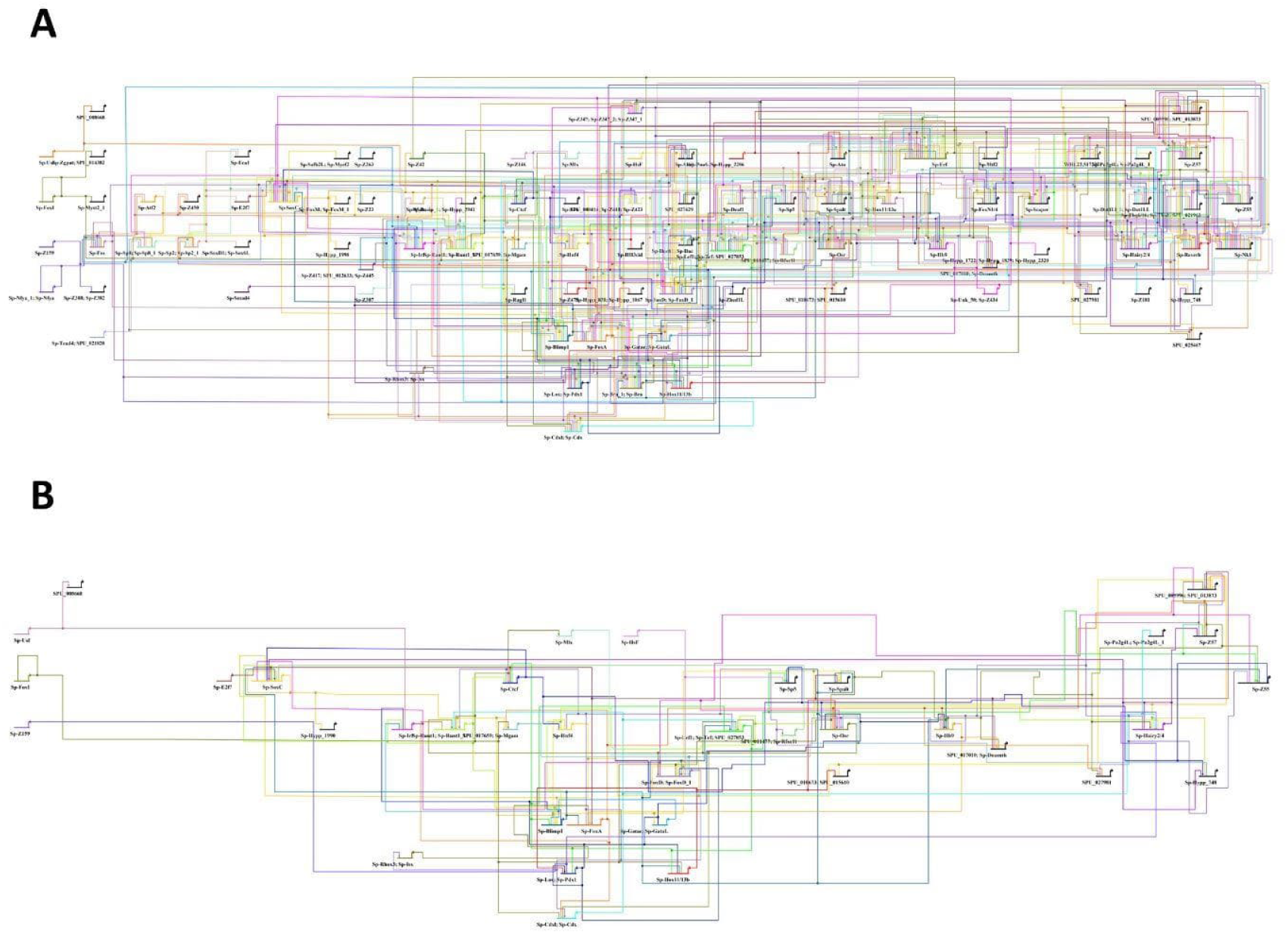
2 dpf *S. purpuratus* gastrula stage hindgut gene regulatory networks. **A)** Full gene regulatory network of the transcription factors of the 2 dpf sea urchin gastrula hindgut (1) cluster. **B)** Full gene regulatory network of the transcription factors of the Sp-Pdx1 positive cells within the hindgut (1) cluster.

**Supplementary Figure 4.**
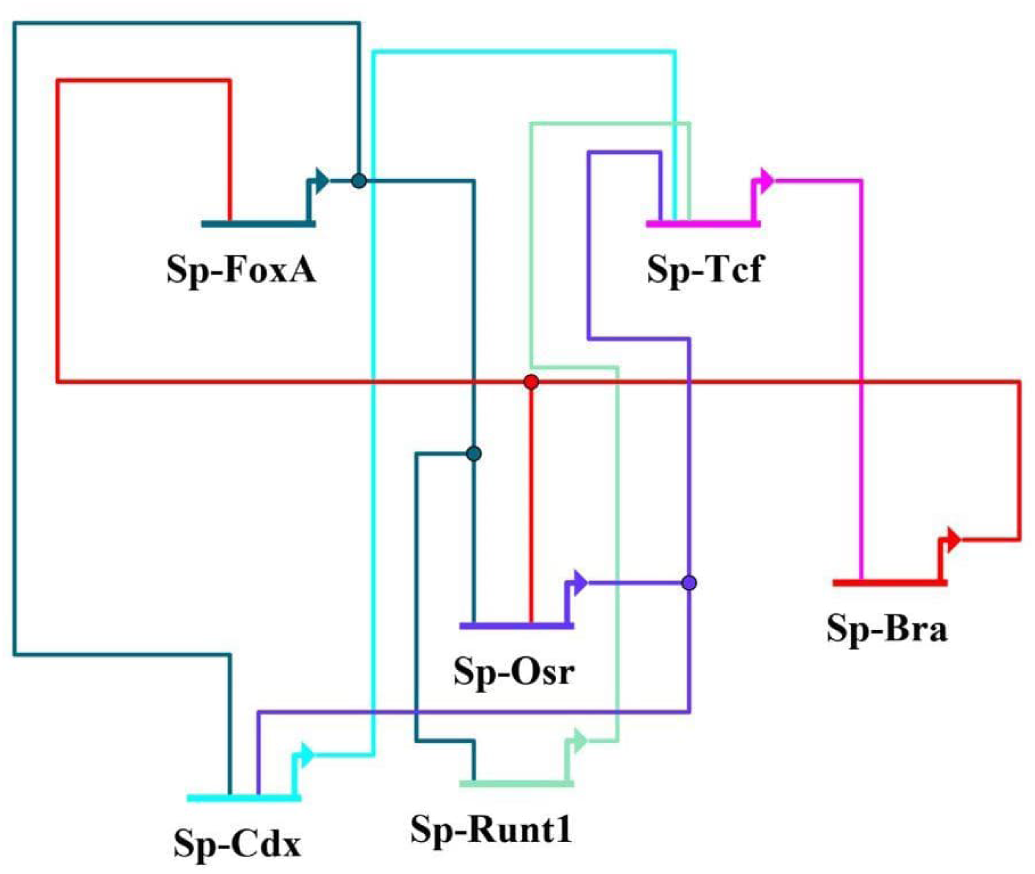
Gene regulatory network connecting Sp-FoxA and Sp-Bra within the published GRN highlighting the indirect nature of this interaction.

**Supplementary Figure 5.**
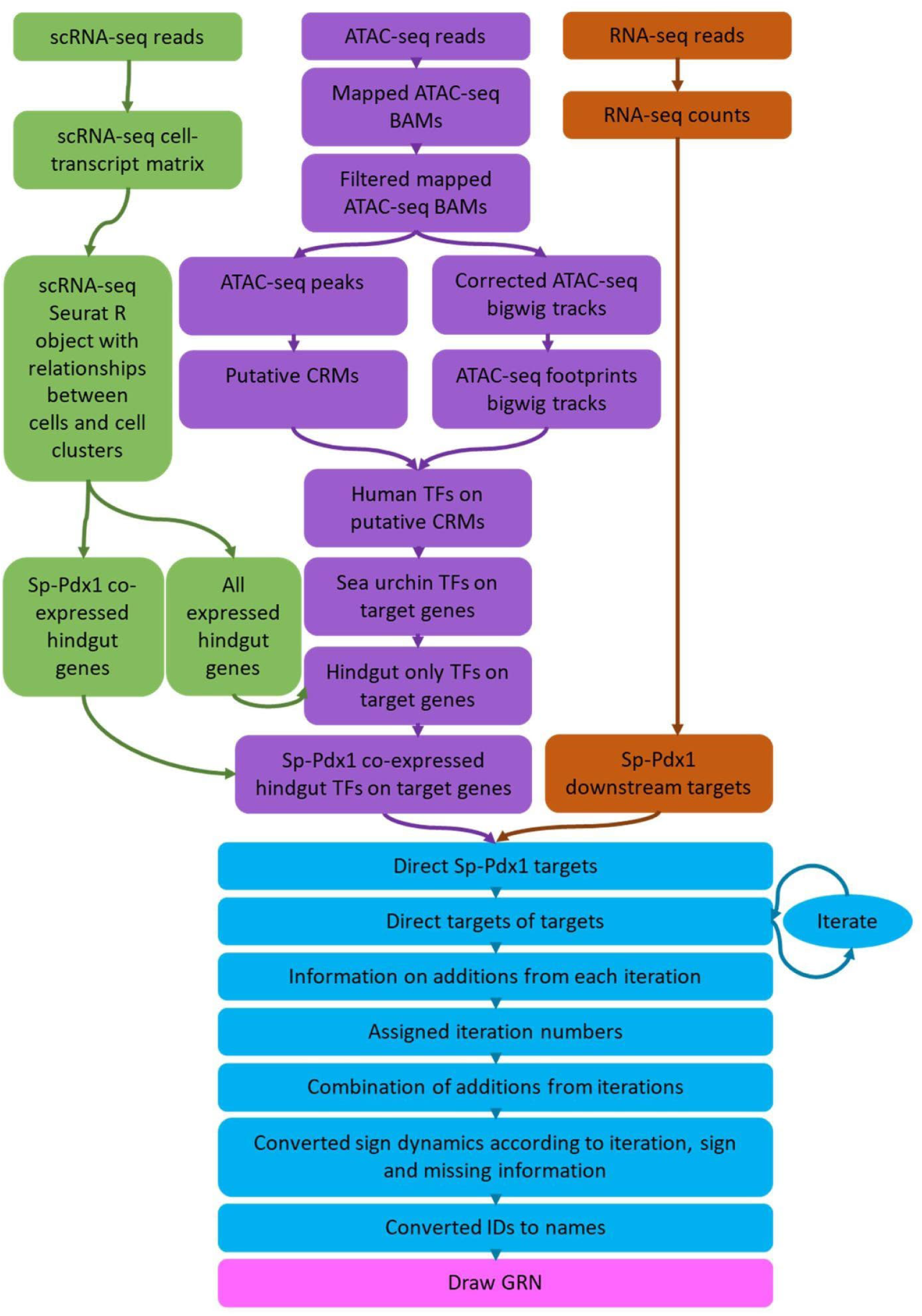
Flowchart of the *in silico* GRN drafting pipeline. Colors represent analyses performed on different data types. Analysis of scRNA-seq data is green, analysis of ATAC-seq data is purple, analysis of differential RNA-seq following Sp-Pdx1 knockdown is brown, combination of the datasets and downstream analysis is blue, drawing the GRN is pink.

**Supplementary Figure 6.**
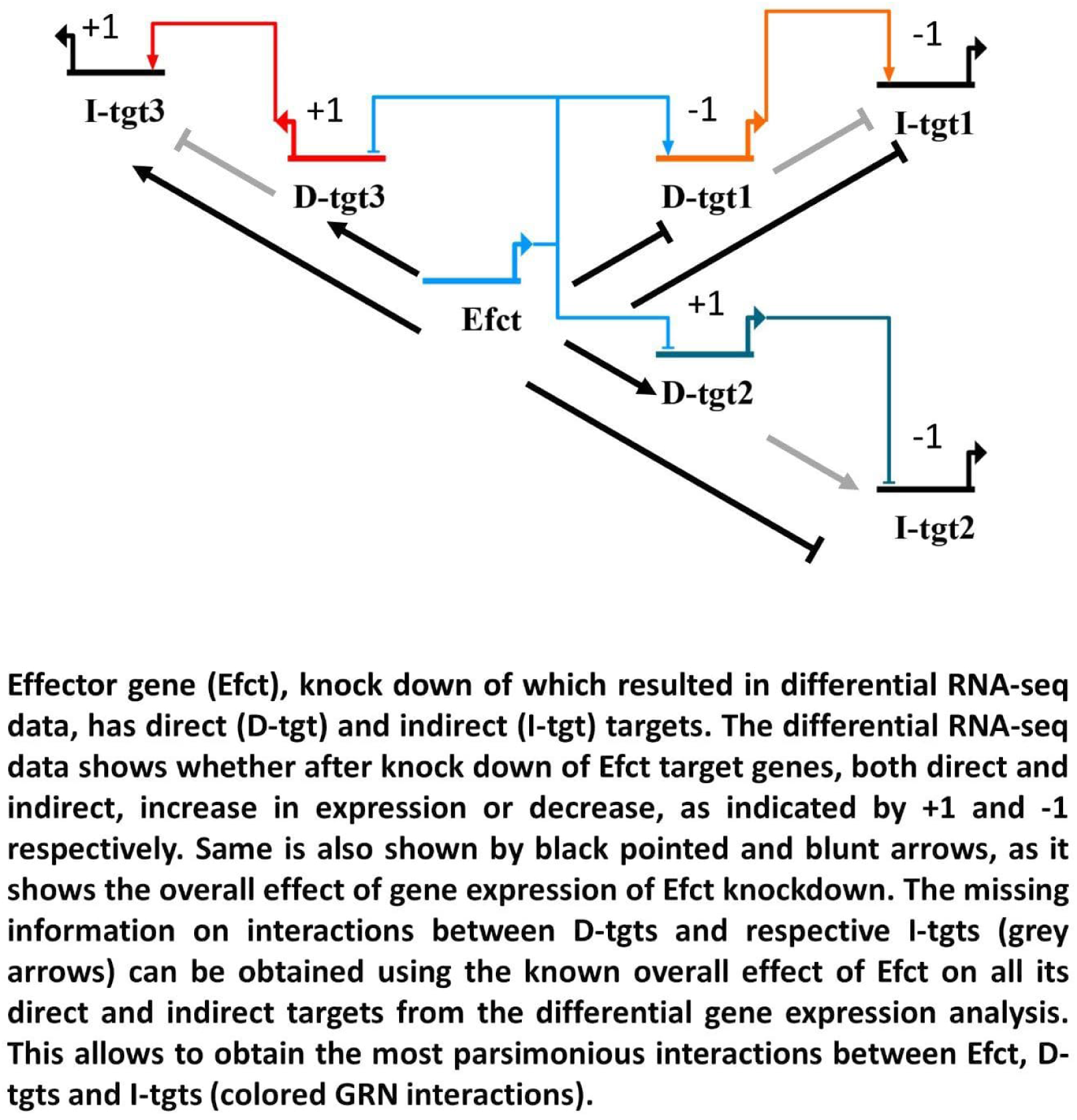
Schematic representation of sign assignment logic used for GRN drafting.

